# Exploring a pico-well based scRNA-seq method (HIVE) for simplified processing of equine bronchoalveolar lavage cells

**DOI:** 10.1101/2024.11.02.621659

**Authors:** Kim Fegraeus, Miia Riihimäki, Jessica Nordlund, Srinivas Akula, Sara Wernersson, Amanda Raine

## Abstract

Single-cell RNA sequencing (scRNA-seq) is a valuable tool for investigating cellular heterogeneity in diseases such as equine asthma (EA). This study evaluates the HIVE™ scRNA-seq method, a pico-well-based technology, for processing bronchoalveolar lavage (BAL) cells from horses with EA. The HIVE method offers practical advantages, including compatibility with both field and clinical settings, as well as a gentle workflow suited for handling sensitive cells.

Our results show that the major cell types in equine BAL were successfully identified; however, the proportions of T cells and macrophages deviated from cytological expectations, with macrophages being overrepresented and T cells underrepresented. Despite these limitations, the HIVE method confirmed previously identified T cell and macrophage subpopulations and defined other BAL cell subsets. However, compared to previous studies T helper subsets were less clearly defined.

Additionally, consistent with previous scRNA-seq studies, the HIVE method detected fewer granulocytes and mast cells than anticipated in the total BAL samples. Nevertheless, applying the method to purified mast cells recovered an expected number of cells. A small set of eosinophils were also detected which have not been characterized in earlier studies. In summary these findings suggest that while the HIVE method shows promise for certain applications, further optimization is needed to improve the accuracy of cell type representation, particularly for granulocytes and mast cells, in BAL samples.

## Introduction

Single-cell RNA sequencing (scRNA-seq) has quickly become a powerful tool for analyzing the transcriptomes of thousands of individual cells across various tissues and organisms. This technology has provided unprecedented insights into cellular function and diversity, both in health and disease. Although scRNA-seq is widely used in human research, there remains a relative lack of similar studies in equines (1–4)

Horses are a useful model for studying a variety of diseases, including allergies, autoimmune disorders, and respiratory conditions (5–8). Along with humans and cats, horses are one of the few mammals that naturally develop asthma, making them an important model for studying this condition. Equine asthma (EA), similar to human asthma, is a complex and heterogeneous disease that affects horses of different breeds and ages. EA significantly compromises the welfare of affected horses and is a prevalent cause of suboptimal performance in sport horses (9). The clinical symptoms of EA include decreased performance, coughing, increased respiratory effort, and mucus accumulation. EA is categorized into two subtypes: mild-moderate asthma (mEA) and severe asthma (sEA) (10–12).

A key characteristic of all forms of EA is the increased infiltration of specific immune cells, such as neutrophils, mast cells and occasionally eosinophils, into the lungs (13,14). This leads to higher proportions of these cells in bronchoalveolar lavage (BAL) samples from affected horses. Given their role in EA pathogenesis, studying these cells in greater detail— especially in the context of other immune cell types present in BAL—is essential. However, both granulocytes (neutrophils, eosinophils) and mast cells are highly sensitive (15,16), partly due to their high RNase activity, which requires careful handling to prevent cellular stress and cell death.

Previous scRNA-seq studies have investigated gene expression differences in specific BAL cell types between asthmatic and healthy horses using droplet microfluidics (1,2). While these studies successfully identified major BAL cell types, they reported low recovery rates for granulocytes and mast cells. It remains unclear whether these low recovery rates were due to the microfluidic cell partitioning methods or suboptimal sample preparation.

Additionally, the standard scRNA-seq sample collection methods can be impractical in field (i.e performing sampling directly at a study site) or clinical settings, where access to specialized equipment is limited. When fresh cells are used, prompt sample handling is ideal, preferably avoiding prolonged storage on ice for sensitive cell types. Cryopreservation or cell fixation techniques are alternatives when immediate processing is not possible; however, these methods require centrifugation and washing steps, often involving swing-bucket rotors, which may not be practical in field settings. These constraints affect certain types of studies, such as those investigating the cellular response to air quality in stable environments and training-associated inflammation in equine airways, where sampling is preferably performed on-site.

This study aimed to evaluate the HIVE™ single-cell RNA-seq solution (Honeycomb Biotechnologies) as an alternative method for processing equine BAL cell samples. The HIVE workflow offers several advantages for studying lung immune cell populations. First, HIVE is designed for sensitive cells, utilizing a gravity-based system that gently captures and stabilizes cells in pico-wells (17). The collector cartridges can be shipped and stored until batch processing. Second, the simple and rapid cell partitioning using portable cartridges allows scRNA-seq sampling in field and clinical settings with only basic equipment, like pipettes. Third, the HIVE system accommodates diluted cell suspensions and large volumes, making it particularly useful for BAL sample studies by eliminating the need for centrifugation, thereby reducing cell stress.

To assess the HIVE system’s performance, we analyzed BAL samples from horses undergoing clinical evaluation for EA. The resulting transcriptomic profiles of major cell types were comparable to those obtained from a previous study using the Drop-seq method. Sub-clustering revealed both previously identified populations and novel cell subsets. However, the HIVE method was less successful in preserving the cell type compositions determined by cytology compared to previous approaches.

## Material and methods

### Study design and sample collection

BAL cells were sampled from horses undergoing clinical evaluation for EA at the University Animal Hospital, Swedish University of Agricultural Sciences (SLU). The procedure was performed as part of routine diagnostic practice due to clinical signs of EA. The examination and BAL sampling were carried out by the same veterinarian for all horses, following the procedure previously outlined by Riihimäki et al (1). To optimize cell viability for scRNA-seq, the cells were processed for transcriptomic analysis before receiving any results from cytology analysis. Descriptive information for all horses (age, sex, breed and sampling dates are provided in S1. Table. Clinical information has been deposited at the figshare repository associated with this study. The samples were collected at convenience and there were no specific inclusion criteria applied. The study was approved by the Uppsala regional ethical review board (5.8.18-20690/2020) and all horse owners approved by written consent. For the Drop-seq samples that were re-analyzed in this study, clinical information has been previously published (1).

### Cell capture, library preparation and sequencing

After sample collection, the BAL samples were kept at 4°C, transported to the lab on ice and processed within 2-4 hours of sampling. Cytospin analysis was performed as previously described (1). Cell concentrations and viability were determined with the Cellometer K2 automated cell counter (Nexcelom Bioscience, Lawrence, MA, USA). The viability of the BAL cells was measured to be > 90 % by the Nexcelom Matrix software (AO/PI staining), for all samples included in the study. Cells were then isolated and stabilized in HIVE pico-wells, following the manufacturer’s instructions for the HIVE v.1 Sample capture kit (Honeycomb Biotechnologies). There are two options to apply cells to the HIVE collectors: i) by gravity flow or ii) by centrifugation. The first option was used in this study. Briefly, in order to remove debris, BAL solutions were passed through a cell strainer (70 μM), according to standard practice procedures for scRNA-sampling. As a result, cells trapped in mucus may have been lost during cell straining. A volume containing 15,000-30,000 cells (35-85 μl) was diluted in 1 ml PBS + 0.01 % BSA, just prior to loading the HIVE collector. The cell suspension (1 ml) was applied to the sample collector and incubated for 30 minutes at room temperature. The collectors were then washed with 1 ml of Wash Buffer before 1 ml of Cell Recovery Solution was added and stored at −20 °C until further processing. For two of the horses (KA & QU), different cell numbers were applied to two HIVE collectors each (Table 1). Subsequent library preparation was performed according to the HIVE single-cell RNA-seq Processing Kit v1 protocol using 9 PCR cycles in the Index PCR step. The size profiles of the amplified sequencing libraries were determined using TapeStation and the final library concentrations were measured with qPCR. Examples of TapeStation library traces are shown in Supplementary Figure 1. The libraries were sequenced on an Illumina NovaSeq6000 system using the custom sequencing primers included in the HIVE library preparation kit and a 28+8+8+90 read set up and aiming at between 50-100K reads/cell.

**Table 1.** Cytology data for the BAL samples analyzed with the HIVE method.

| Horse #<br>and ID | HIVE<br>library # | Leukocytes<br>(10e6/L) | Neutrophils<br>(%) | Lymphocytes<br>(%) | Macrophages<br>(%) | Eosinophils<br>(%) | Mast<br>cells<br>(%) |
| --- | --- | --- | --- | --- | --- | --- | --- |
| 1 NA | 1 | 280 | 1 | 42 | 55 | 1 | 2 |
| 2 DE | 2 | 280 | 1 | 24 | 65 | 0 | 10 |
| 3 QU | 3,4 | 110 | 0 | 30 | 61 | 0 | 9 |
| 4 AM | 5 | 260 | 4 | 34 | 58 | 0 | 4 |
| **5 ST** | 6 | 180 | 4 | 29 | 53 | 0 | 14 |
| **6 BE** | 7 | 170 | 0 | 42 | 51 | 1 | 6 |
| **7 FA** | 8 | 410 | 1 | 21 | 66 | 0 | 11 |
| **8 LE** | 9 | 240 | 3 | 18 | 72 | 3 | 4 |
| **9 KA** | 10,11 | 400 | 1 | 3 | 72 | 5 | 19 |
| **10 GN** | 12 | 430 | 31 | 25 | 41 | 0 | 3 |
| **11 FL** | 13 | 360 | 3 | 39 | 56 | 0 | 3 |
| **12 UF\*** | 14 | 370 | 4 | 26 | 65 | 2 | 3 |
| **13 AT^** | 15,16 | 340 | 3 | 38 | 54 | 1 | 4 |
| **14 TT^** | 17,18 | 400 | 7 | 26 | 65 | 0 | 2 |

\*Purified mast cells. ^ Experiment with alternative sample handling conditions.

### Purification of mast cells

Mast cells were isolated from a single equine BAL sample using the MACSQuant Tyto cell sorter, as detailed in *Akula* et al (18). The total procedure took 5 hrs. from the time of BAL sampling. Mast cells were kept on ice at all times. The purity of the BALF MCs was estimated to be 92.9% (18). An aliquot of the purified mast cells from the BAL sample described in that study (denoted MC1 in *Akula* el, herein denoted by the horse ID=UF) was analyzed with HIVE. Immediately after the purification procedure 15,000 purified mast cells were diluted in 1 ml cold PBS + 0.01 % BSA and applied to the HIVE collector. Libraries were then prepared, sequenced and analyzed following the same procedure as stated above.

### Test of alternative cell sample handling conditions

BAL samples (AT and TT, Table 1) were immediately split into two aliquots directly after endoscopy and BAL collection. One aliquot was kept on ice until HIVE loading, while the other aliquot was kept at room temperature. Just before applying the cells to the collectors, they were diluted in 1 ml of RPMI media (cold or room tempered, respectively) containing 5% FBS, and 10 μl of RiboLock RNase inhibitor. The duration from BAL sampling to loading the cells onto the HIVE collectors was less than one hour.

### Data pre-processing

Raw sequencing data (fastq files) were converted to count matrices of gene expression values using the custom software BeeNet^TM^ (Honeycomb Biotechnologies). A custom Python script was used to extend the annotations for the 3’-untranslated regions of the genes in the reference genome (Equus caballus, NCBI annotation release 103) with 1000 bp (except for when the extension overlapped a neighboring gene). Following the recommendations in the HIVE scRNAseq BeeNet^TM^ (v.1.1) software guide, the number of expected barcodes in each sample was set to 40% of the starting input cell number (6,000-12,000 cells per sample, Table 1). BeeNet output metrics are listed in S1 Data.

### Quality control and initial clustering analysis of scRNA-seq libraries

Quality control (QC) and further downstream analyses were performed using the R package Seurat (v 4.3.0.1) (19). For the initial HIVE libraries cellular barcodes with < 350 detected genes, >10% mitochondrial reads, >15% ribosomal reads and/or >8,000 UMI counts were filtered out. Doublet detection was performed using the DoubletFinder_v3 (20) package for each sample. After filtering, a total of 55,759 cells remained for downstream analysis. Normalization and scaling were performed using the SCTransform function (v. 0.3.5) with 3,000 variable features, regressing out the number of features/cells. Integration was then performed, on sample basis, with Seurats RPCA method (k.anchor = 20).

The CellCycleScoring function in the Seurat package was used to investigate if the cell cycle stage had an effect on clustering. Following the same procedure as Sage et al^1^ the Biomart package was used to convert the lists of human markers for the G2M and S phase (“cc.genes.updated.2019” from Seurat) to their equine orthologs. Based on the scores obtained, cells were then visualized according to G2M phase (cycling) or S phase (resting). Dimensionality reduction was performed using principal component analysis (PCA) and the number of principal components (PCs) were set to 25. The number of PCs were determined by inspection of scree plots and computation of the number of PCs exhibiting > 0.1 % variation. Clustering was performed using the FindNeighbors function in Seurat, with the Louvain algorithm as default. The FindClusters function was then used, and clustering resolutions between 0.1 and 1 were assessed. Based on visual inspection of the various clustering resolutions (UMAP plots) as well as cluster stability (assessed with the clustree function (21), and assessment of cluster specific gene signatures, a clustering resolution of 0.3 was selected for the initial clustering of the total BAL population. Clusters with similar gene expressions were collapsed into the major cell-types when computing cell-type specific markers. Statistical inference of differences in cell type composition as compared with cytology was computed using the Wilcoxon signed-rank test in R.

The four HIVE libraries used to investigate alternative sampling conditions were integrated and analyzed as described above (n = 10,000 cells, clust res 0.2, 10 PCs). The isolated mast cells were also analyzed using the same filtering and clustering steps as described above, except for the number of detected genes, where the limit was set to 275 (n= 2,140 cells, 10 PCs, clust res 0.2).

Noteworthy, mast cell tryptase genes appear to be absent from the current equine reference annotation. However, comparison of the protein sequences of human mast cell-specific tryptases and the protein sequence of the equine RET gene (NP_001075344.1, NP_003285.2, NP_077078.5) reveal high homology, including conservation of the catalytic triad (22), strongly suggesting that equine TPSAB1/TPSB2 is misannotated as RET.

### Comparison of mastocytic HIVE and Drop-seq samples

HIVE and previously generated DropSeq data from BAL samples with more than 3% mast cells (based on cytology) were selected for re-analysis and comparison. This included 11 HIVE libraries and 8 DropSeq libraries. The HIVE data was processed as described above, while the DropSeq data was obtained from Riihimäki et al (1). The EmptyDrops (23) method had previously been applied to remove empty droplets from those count matrices, allowing for a lower threshold to filter out low-quality cells (<200 genes per cell). Similarly, for the HIVE data, cells with more than 10% mitochondrial reads, more than 15% ribosomal reads, and/or more than 8,000 UMIs were excluded. The mastocytic HIVE and Drop-Seq datasets were analyzed separately using the same computational approach for doublet removal, integration, and clustering. The number of principal components (PCs) was set to 20, and a cluster resolution of 0.1 was selected for both datasets to visualize major cell types.

### Cluster annotation and independent re-clustering of cell types

To identify genes differentially expressed between clusters, the PrepSCTFindMarkers and FindAllMarkers functions in Seurat were used, using an average log2fold change (log2FC) threshold of >0.2 and adjusted P-value < 0.05. These markers, in combination with canonical markers previously used in horse and human scRNA-seq studies (1–4,24), were used to annotate the cell clusters into the major cell types. After identifying the major cell types, the Subset function in Seurat was used for sub-clustering of each cell type; alveolar macrophages (AMs), T cells, neutrophils, mast cells and dendritic cells (DCs), following the same procedure as described above, but with different number of PCs and clustering resolutions (AMs: 15 PCs, clustering resolution of 0.4, T cells: 12 PCs, clustering resolution of 0.6, mast cells: 5 PCs, clustering resolution of 0.2, neutrophils: 10 PCs, clustering resolution of 0.2, dendritic cells: 10 PCs, clustering resolution of 0.1). A population of cells exhibiting expression of both T cell and macrophage genes (3000 cells, 5 % of total BAL cells) were removed when re-clustering the T cells.

## Results and Discussion

### Initial evaluation of the HIVE method for clinical BAL samples

Clinical BAL samples from eleven horses undergoing evaluation for EA were initially analyzed with the HIVE v.1 solution (Honeycomb Biotechnologies) (Figure 1). Healthy controls were not included in this method evaluation as one of the main objectives was to investigate whether the HIVE method could improve the recovery of neutrophils and mast cells, which are typically present in low numbers in normal BAL samples. Thus, the study did not aim to conduct research on EA but rather to explore the suitability of the HIVE method for processing BAL samples for future EA studies. Duplicate libraries with different numbers of cell loadings were analyzed for two of the BAL samples, so that in total 13 HIVE libraries were analyzed in the initial round. In ten out of eleven of the BAL samples, cytology indicated ongoing airway inflammation (3% ≥ mast cells and/or eosinophils or 10 % ≥ neutrophils). The majority of horses exhibited mastocytic inflammation, although one horse had severe neutrophilic inflammation. Total cell counts in the BAL were between 100-400 cells/ul which meant that an appropriate number of cells for loading the sample collector (15-30,000 cells) could easily be obtained by diluting a small volume from the original BAL sample directly into 1 ml of loading buffer, limiting stress to cells by centrifugation. The cell compositions of the BAL samples, as assessed by cytology staining, are shown in Table 1.

**Figure 1.**
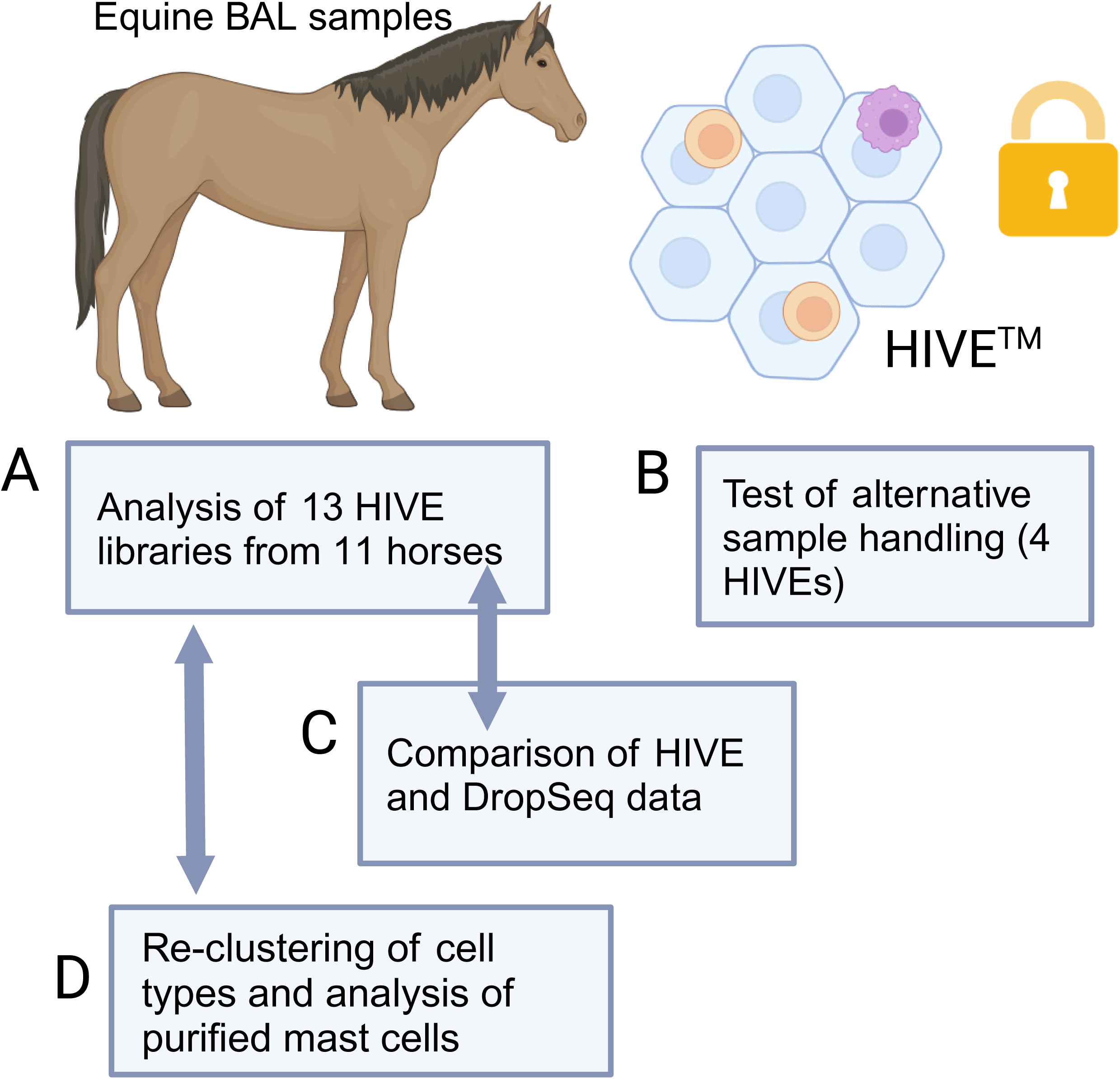
Overall study outline. A) Thirteen libraries obtained from eleven equine BAL samples were initially prepared and analyzed using HIVE (v.1). B) Four HIVE libraries (from two additional BAL samples) were prepared with alternative sample buffers and storage temperatures to investigate whether cell type recovery would be improved by optimizing sample handling conditions. C) Since it was not possible to compare HIVE data from the same samples with an alternative method, HIVE data was compared to another method (Drop-seq) using samples that exhibited similar BAL phenotypes (mastocytic). D) The major BAL cell subsets from the initial thirteen HIVE libraries were independently sub-clustered. Additionally, a HIVE library prepared from mast cells purified from an additional BAL sample was analyzed. Created in BioRender. Raine, A. (2024) BioRender.com/n54f847

After QC and removal of doublet cells, a total of 55,759 cells remained for the downstream analysis. The final number of cells recovered per sample varied from 1,481 to 7,358 (Table 2). A median of 11,666 genes/sample were detected (counting genes detected in ≥ 10 cells). The median number of genes/cells varied between 492-1,189 across the individual samples. The mean percentage of mitochondrial and ribosomal reads (across all samples) were 2 and 6 %, respectively, which is not higher than what has been reported for other methods (25). Violin plots illustrating the common metrics for assessing the quality of the data are presented in S2 Figure and sequencing/mapping metrics in S1 Data).

**Table 2.**
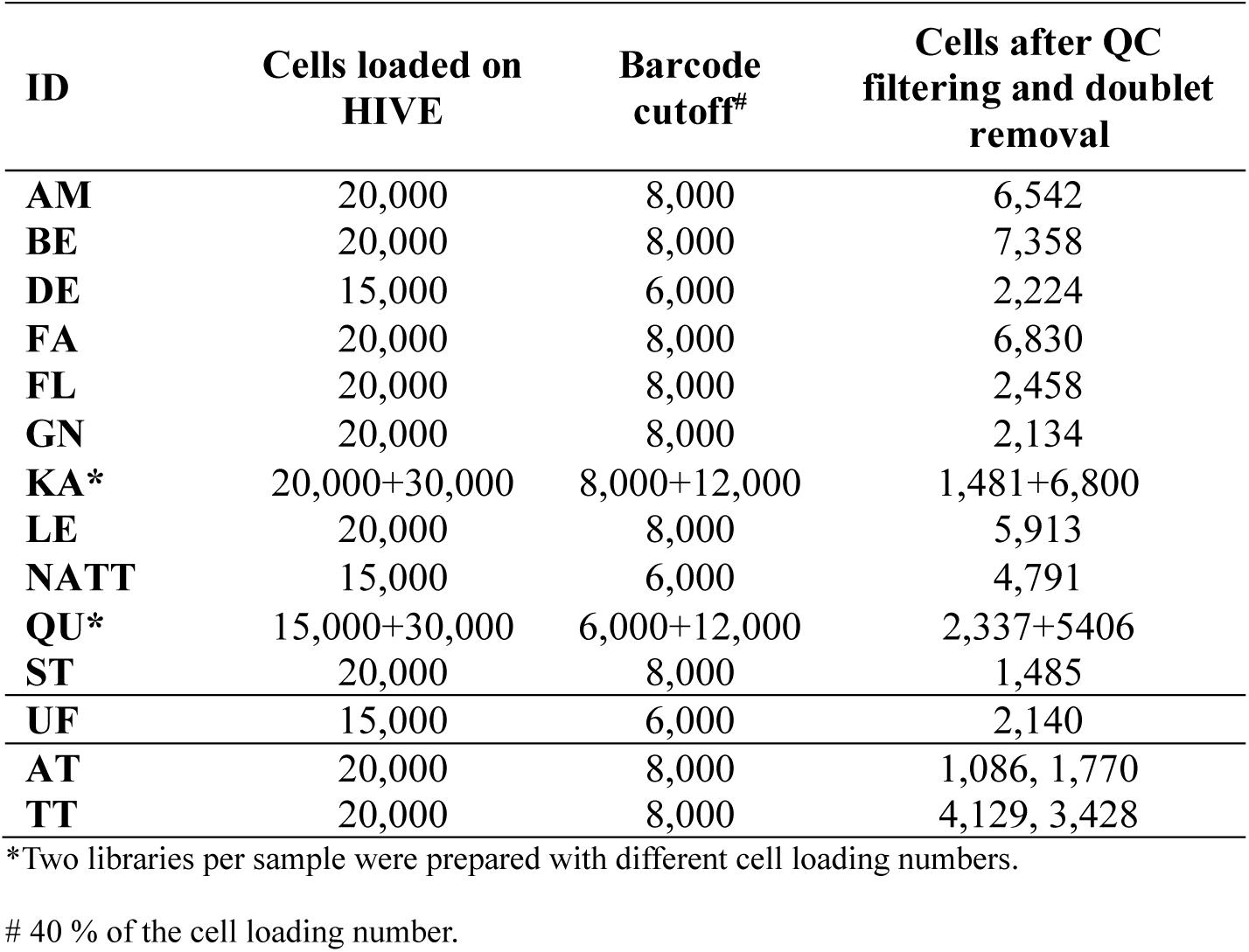
Number of cells loaded per HIVE collector and number of cells recapitulated after QC filtering.

The HIVE data recovered the expected five major cell types in BAL and their gene expression signatures were essentially similar to those observed in previous equine scRNA-studies: alveolar macrophages (n = 39,800, top cell type markers = *WFDC2*, *PSAP*, *APOE*, *CD163*), proliferating macrophages (n = 2,149, *CENPF, TOP2A*), T cells (n = 10,129, *RESF, STK17B*, *GZMA*, *CD3E*), mast cells (n = 737, *MS4A2, FCER1A, GNPTAB*), neutrophils (n = 1487, *SNX10*, *GBP5*, *CXCL1*, *CXCL8*) and dendritic cells (n = 1187, *DRA*, *CST3*, *CD74*) (Figure 2 and S2 Data). However, cell-type proportions in the scRNA-seq data differed considerably compared to cytology counts (Figure 2). Low numbers of neutrophils (P value = 0.3, not significant) and mast cells (P value < 0.001, Wilcoxon signed-rank test) were detected, even in samples that exhibited very high proportions of these cells according to cytology. Moreover, in the HIVE samples, an average 74 % of the cells were annotated as macrophages, which was higher than the counts observed in cytology (mean 59 %, P value < 0.001, Wilcoxon signed-rank test). For T cells, the percentages were lower than expected compared to cytology (mean: 19% vs. 28% in cytology, P value < 0.001). Notably, cytology indicated a very low number of lymphocytes (3%) in one horse (KA), while the corresponding number was 10% in the two HIVE libraries from that horse.

**Figure 2.**
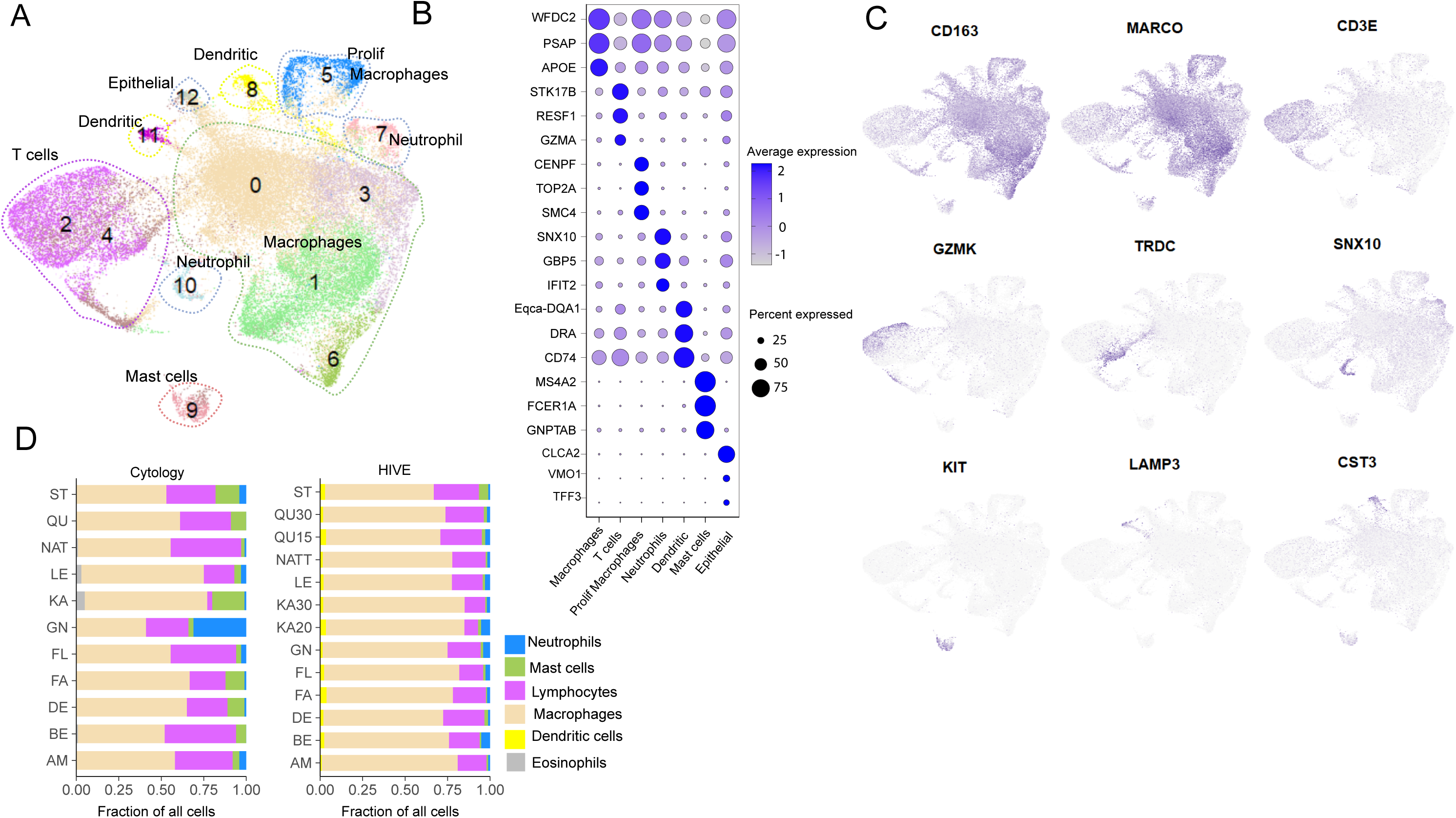
scRNA-seq generated from equine BAL samples using the HIVE method. A) BAL samples from eleven horses undergoing evaluation for equine asthma were analyzed using the HIVE pico-well technology. Duplicate libraries with different cell loadings were prepared from two horses (KA & QU), thus a total of thirteen HIVE libraries were analyzed. Cluster representations of integrated HIVE data are shown as UMAP plots. The expected cell types in equine BAL were identified and clusters labelled accordingly: alveolar macrophages (clusters 0, 1, 3, and 6), proliferating macrophages (cluster 5), T cells (clusters 2 and 4), neutrophils (clusters 7 and 10), mast cells (cluster 9), dendritic cells (clusters 8 and 11), and a small population of epithelial cells (cluster 12). B) Bubble plots showing the top expressed marker genes after collapsing clusters into major cell types. C) Feature plots visualizing the expression of a selection of cell-type specific marker genes on the UMAP. D) Cell type proportions computed from HIVE scRNA-seq data versus cytology data, per horse. Macrophage numbers were significantly higher, whereas T cell, neutrophil, and mast cell numbers were lower in HIVE data compared with cytological expectations.

### A preliminary investigation into the impact of sample-handling conditions on BAL cell-type recapitulation in HIVE data

To explore potential improvements in cell type recovery through modifications in sample processing conditions—such as temperature, cell dilution buffer, and handling time —we generated four additional HIVE libraries from two additional horses (Horse AT and TT; HIVE libraries #15-18, as shown in Table 1). The addition of RNase inhibitors to buffers and avoiding storage on ice has previously been recommended for neutrophil sample preparation. (https://kb.10xgenomics.com/hc/en-us/articles/360004024032-Can-I-process-neutrophils-or-other-granulocytes-using-10x-Single-Cell-applications). In this preliminary experiment, all three aforementioned factors were modified simultaneously because high costs prevented a more systematic approach.

After the BAL procedure, each sample was promptly split into two aliquots: one kept at room temperature and the other on ice. Cells were diluted in an alternative buffer (supplemented with FBS and RNase inhibitor) immediately before loading. Additionally, HIVE loading was completed within one hour, compared to the 2–4-hour window used in earlier tests. An integrated analysis of 10,000 cells from the four libraries suggested that the proportions of T cells, macrophages, and neutrophils were in better agreement with cytology compared to the initial 13 libraries, which were prepared in cold conditions using standard buffer (PBS + 0.01% BSA). However, mast cell recovery remained lower than expected, and eosinophils were not detected (Figure 3, Table 1, S3 Data).

**Figure 3.**
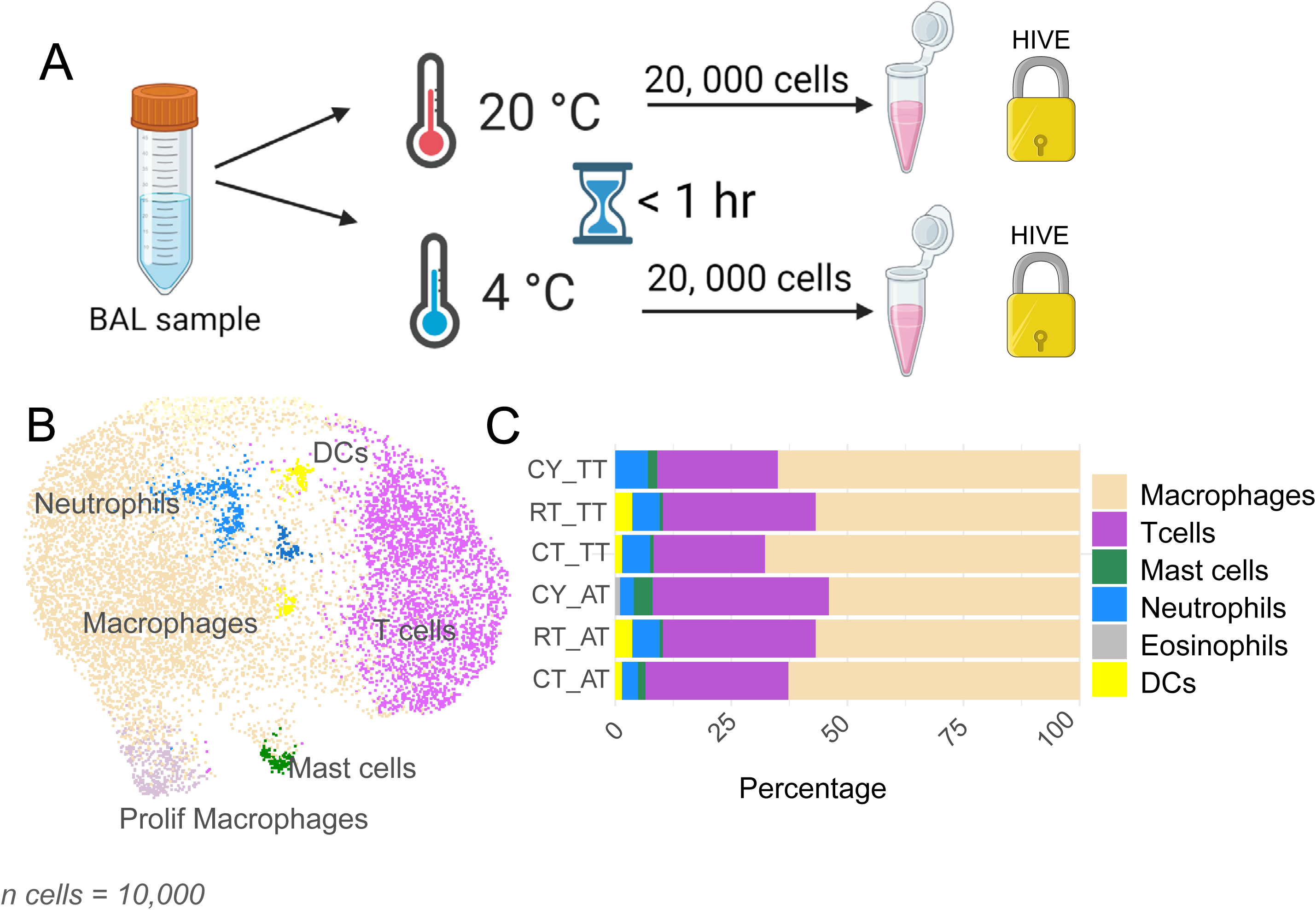
Cell type distribution in HIVE libraries prepared with alternative sample handling conditions. A) Four HIVE libraries were prepared from two BAL samples (IDs: AT & TT) using alternative sample buffers and different sample handling temperatures. Each BAL sample was promptly split into two aliquots and kept either cold or at ambient temperature. Cells were then diluted in cold or ambient temperature cell media buffer (RPMI) supplemented with 5% FBS and RNase inhibitor, directly prior to loading the collectors, which took place in less than one hour after the BAL sampling procedure (as opposed to 2-4 hours in the initial experiments). B) Clustering of integrated data integrated from the four libraries and labelling of the major cell types in BAL. C) Cell type compositions are represented in bar plots and compared to cytology. Although the proportions of macrophages, T cells, and neutrophils appear similar to those in cytology, this interpretation should be made with caution due to the low number of replicates, which precludes statistical testing. Mast cell numbers were still lower in the HIVE data, and eosinophils were not detected at all. CY = cytology, RT = HIVE room temperature, CT = HIVE cold temperature.

The putative improvement in cell type distribution observed for T cells, macrophages and neutrophils should, however, be considered preliminary. Concurrent modification of several experimental variables and the limited number of replicates prevents any meaningful statistical analysis. Therefore, additional studies will be needed to determine the optimal sample handling protocol for equine BAL cells, particularly for samples with high levels of mast cells. One area that warrants further exploration is whether a cell washing step before the HIVE loading, which may reduce RNases, would actually improve the accurate recapitulation of cell types. It should also be noted that previous scRNA-seq studies have successfully analyzed cryopreserved equine BAL cells using 10x Genomics, including the recovery of neutrophils (2,3).

### Comparison of HIVE and Drop-seq data

Next, we compared gene expression profiles obtained from HIVE data with those generated by an alternative scRNA-seq method (Drop-seq) (1). Ideally, method performance would be assessed by analyzing data from the same samples across different protocols. However, the high cost of scRNA-seq and the logistical challenges of processing freshly sampled cells using two different methods made this approach impractical. As a direct comparison was not feasible, we instead re-analyzed previously generated Drop-seq data from equine BAL cells. The same computational approach used for the HIVE data was applied to assess cell clusters and marker genes. To ensure a meaningful comparison, similar sample types were selected (i.e., mastocytic BAL samples with ≥ 3% mast cells).

Thus, gene expression profiles from HIVE libraries prepared from mastocytic BAL samples in the initial experiment (11 libraries, 9 horses, n = 45,222 cells, Table 1) were compared with previously published Drop-seq libraries from mastocytic horses (8 libraries, n = 31,133 cells, S2 Table). Both datasets revealed similar clusters, corresponding to the major BAL cell types (Figure 4). Moreover, very similar marker genes were identified across these cell types (S4 Data). The shared features within the top 25 cell type markers (by log2FC) are listed in Table 3. In contrast to HIVE, T cell and macrophage composition were more consistent with corresponding cytology for Drop-seq data (P value = 0.93, and 0.52, for macrophages and T cells respectively, Wilcoxon signed-rank test). Neutrophil and mast cell recapitulation were significantly low compared with cytology also for Drop-seq (P =0.05 and 0.01, respectively).

**Figure 4:**
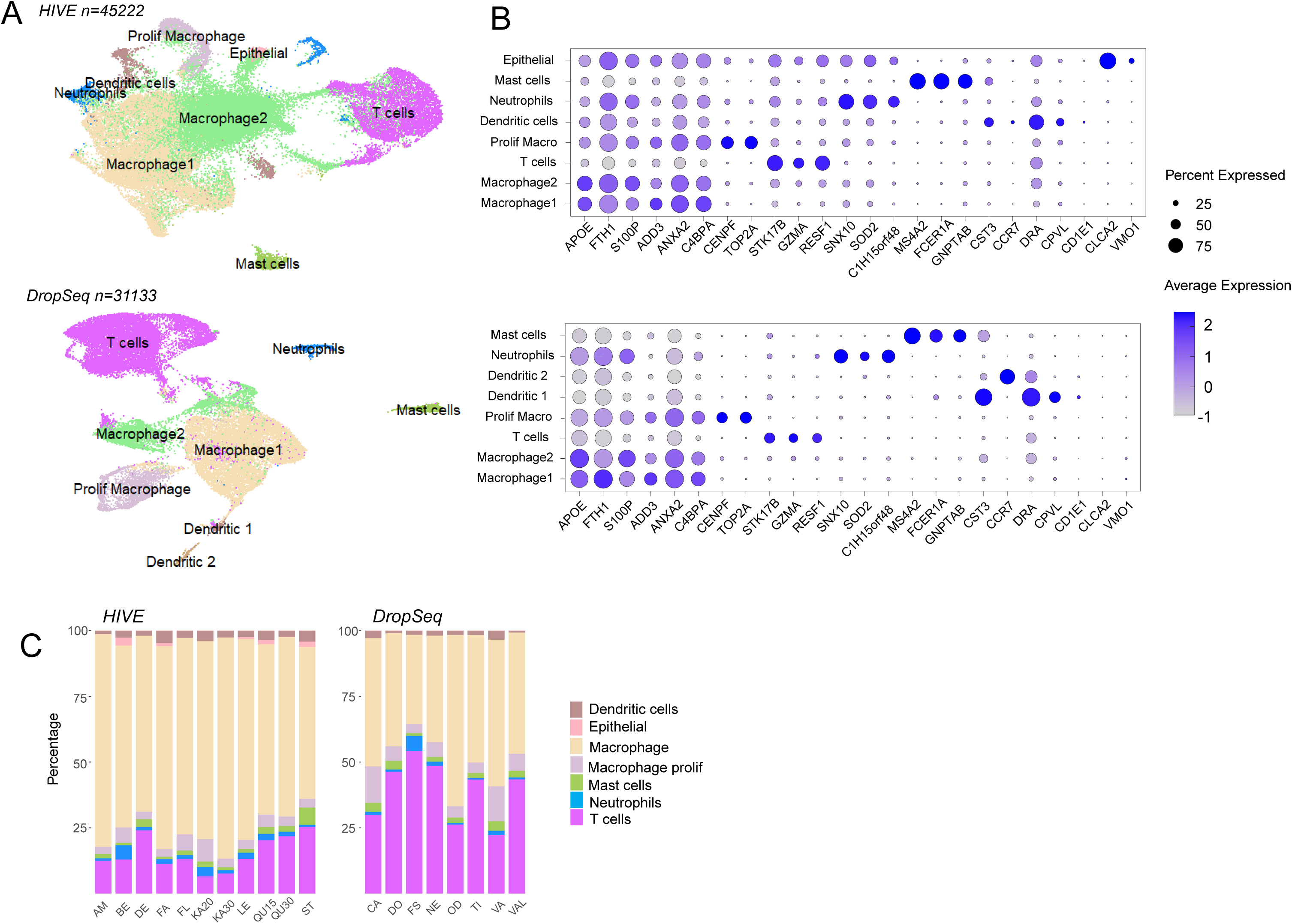
Comparison of clustered HIVE and Drop-seq data from mastocytic BAL samples. A) UMAP representations of HIVE and Drop-seq data. The same major cell types were identified, although epithelial cells were not found in Drop-seq. Epithelial cells are not expected to be present in all BAL samples. B) Bubble plots comparing expression levels of top marker genes across clusters. C) Cell type proportions in individual samples visualized as bar plots. Drop-seq data showed generally higher T cell recovery and thus better agreement with cytology compared to HIVE. Granulocytes and mast cells numbers were similarly low in both data sets.

**Table 3.**
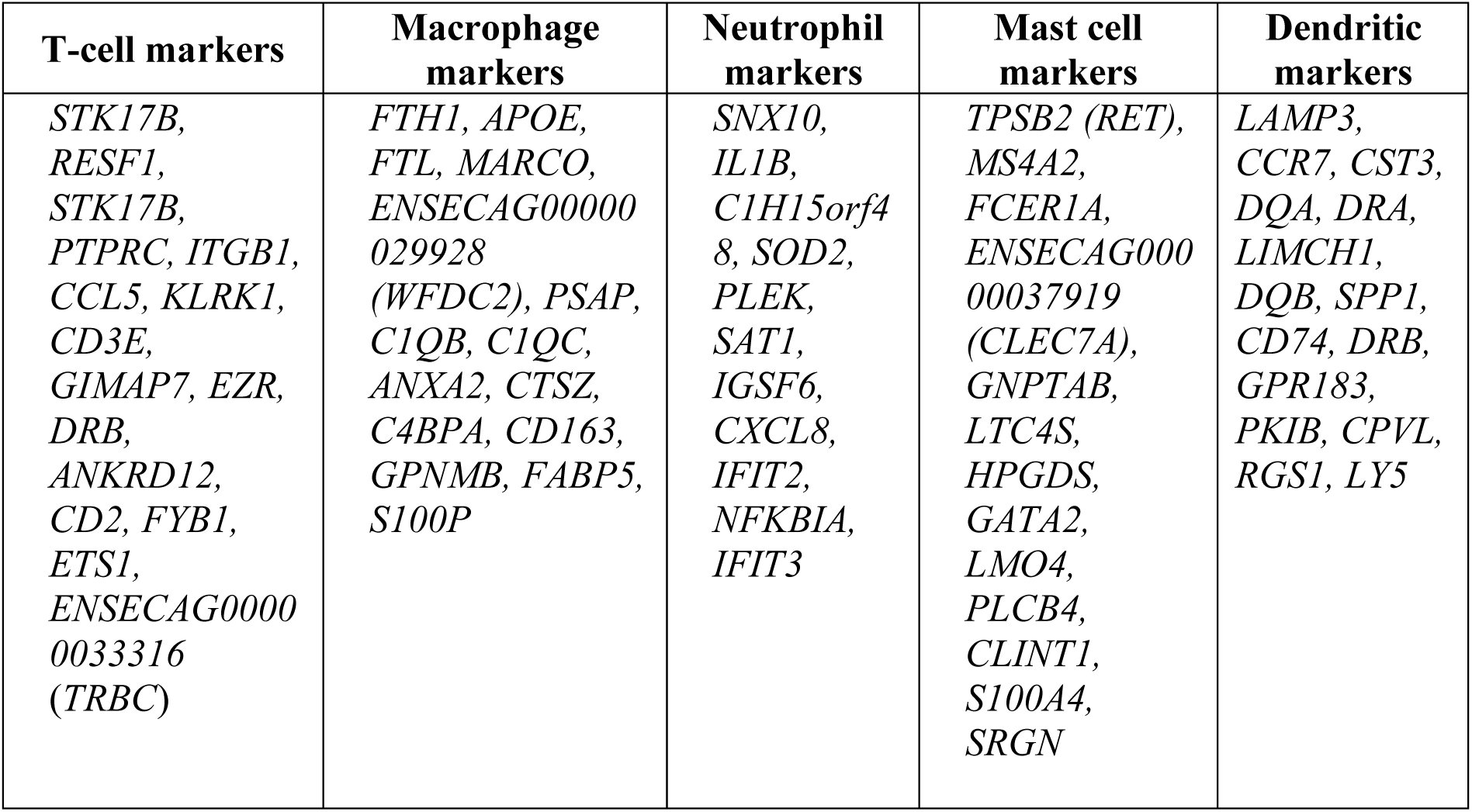
Shared equine BAL cell-type marker genes across HIVE and Drop-seq data. The table lists common cluster markers among the top 25 genes with the highest log2FC in each respective dataset, as identified by Seurat’s (v.4.3) FindAllMarkers function.

In summary, gene expression signatures across the major cell types as identified by both Drop-seq and HIVE were highly similar.

### Independent re-clustering reveal novel and known macrophage and T cell populations in HIVE data

Previous scRNA-seq studies of equine BAL, have identified distinct populations of airway macrophages, along with diverse lung-residing T cell types. To assess whether comparable cell states or types could be identified in the HIVE dataset, independent re-clustering was performed.

#### Macrophages

Given that macrophage polarization occurs along a continuum rather than in discrete stages (26), differences between clusters often manifest subtly in marker expression, with some markers being expressed across multiple clusters. Moreover, the inherent plasticity of macrophages, the limited transcriptomic data available for equine lung-residing macrophages, and technical variabilities in gene detection collectively pose challenges for comparing observed subpopulations across different studies. Nevertheless, re-clustering the HIVE scRNA-seq dataset yielded several clusters which exhibited gene expression signatures akin to previously identified alveolar macrophage (AM) subpopulations (Figure 5 and S5 Data) (1).

**Figure 5.**
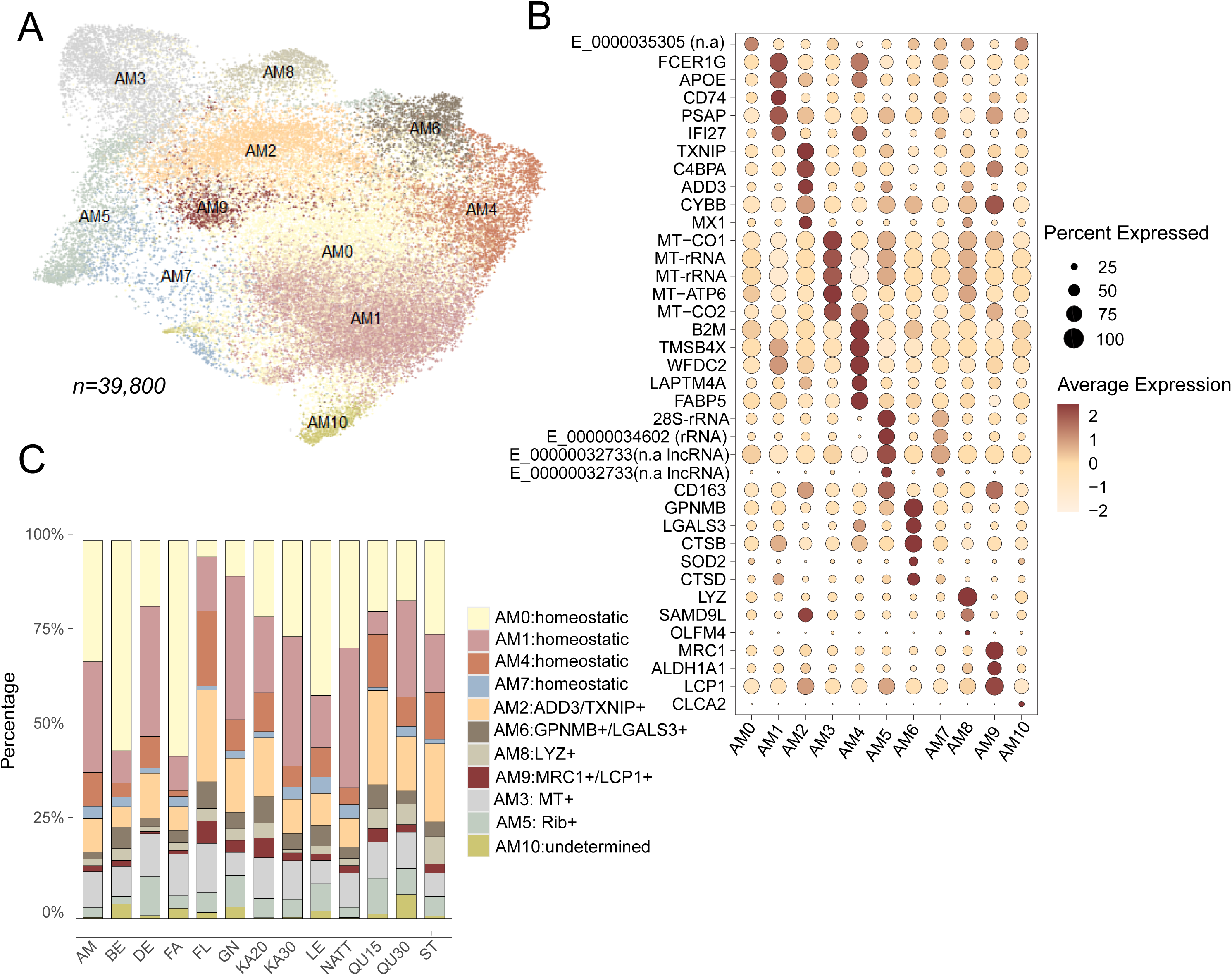
Reclustering of alveolar macrophages. A) Reclustering of 39,800 alveolar macrophages. Clusters AM0, AM1, AM4, and AM7: expression patterns consistent with the homeostatic functions of alveolar macrophages. Cluster AM2: putative activated phenotype (*ADD3+, TXNIP*+). Clusters AM3 and AM5: increased mitochondrial and ribosomal expression, respectively. Cluster AM6: monocyte-derived AMs (*GPNMB+, LGALS3+, MARCO*^low^). Cluster AM8: enhanced expression of lysozyme (*LYZ*+). Cluster AM9: putative anti-inflammatory (*MRC1*+). AM10: undetermined. B) Bubble representation of the top differentially expressed genes across clusters. n.a = gene not annotated in EquCab 3.0. C) Cluster proportions varied across samples, also between libraries prepared from the same BAL samples (KA20, KA30, QU15, QU30). A possible explanation may be is inconsistencies in cluster assignment due to overlapping expression profiles in some AM subsets.

Clusters AM0, AM1, AM4, AM7 exhibited expression profiles consistent with homeostatic functions in alveolar macrophages such as phagocytosis, lipid metabolism (surfactant homeostasis) and immune regulation. Nevertheless, subtle gene expression differences were detected for some of these clusters. For instance, cells in cluster AM1 showed a slight increase in the expression of phagocytic activation-associated genes (*PSAP, APOE*) and genes involved in antigen presentation and immune regulation (*FCER1G, CD74*). AM4 were marked by expression of lipid-associated and cytoskeleton genes (*FABP5, ANXA2, TMSBX4*), immune regulators (*B2M*), protease inhibitors (*WFDC2*), and lysosomal protein (*LAPTM4A*).

Cluster AM2 (*ADD3, TXNIP, CYBB, C4BPA*) may represent a poised or activated state (27), comprising a subset of cells also expressing interferon stimulated (ISG) genes (e.g., *MX1, SAM9DL*). Cluster AM6 (*GPNMB, LGALS3, MARCO*^low^) is suggested to represent monocyte-derived alveolar macrophages (28). Both AM2 and AM6 clusters exhibited expression profiles similar to corresponding populations identified in previous equine scRNA-seq studies (1,2).

Clusters AM3 and AM5 showed upregulation of mitochondrial and ribosomal genes, respectively. Cluster AM8 was characterized by markedly higher expression of lysozyme (*LYZ*). The small cluster AM9 exhibited increased expression of CD206 (*MRC1*), *CD163*, *ALDH1A1*, and *LCP1* suggestive of an anti-inflammatory role (29)

Most individual samples contained cells across all AM clusters, although the proportion of cells in each cluster varied between samples. Notably (Figure 5), there were also some differences in cluster proportions across HIVE libraries from the same BAL sample (KA20, KA30, and QU15, QU30). It remains unclear whether these differences can be attributed to technical bias (in the HIVE method), computational bias (introduced during the clustering of cells that exhibit continuums rather than discrete states), random effects from cell sampling, or a combination of these factors.

#### T cells

Next, independent re-clustering of T cells was performed (Figure 6, S5 Data). A population of cells (∼3,000 cells, 5.5 % of the total cell population) exhibiting expression of both T cell and macrophage genes was identified during the re-clustering of the T cells. These cells appeared scattered throughout the UMAP, though they were assigned to the same cluster. It is unclear whether they represent technical doublets (that DoubletFinder failed to identify), true cell complexes or a combination of both. However, as this cluster exhibited ambiguous gene signatures (which interfered with clustering) it was removed and the remaining T cells (n= 6,672) were sub-clustered.

**Figure 6.**
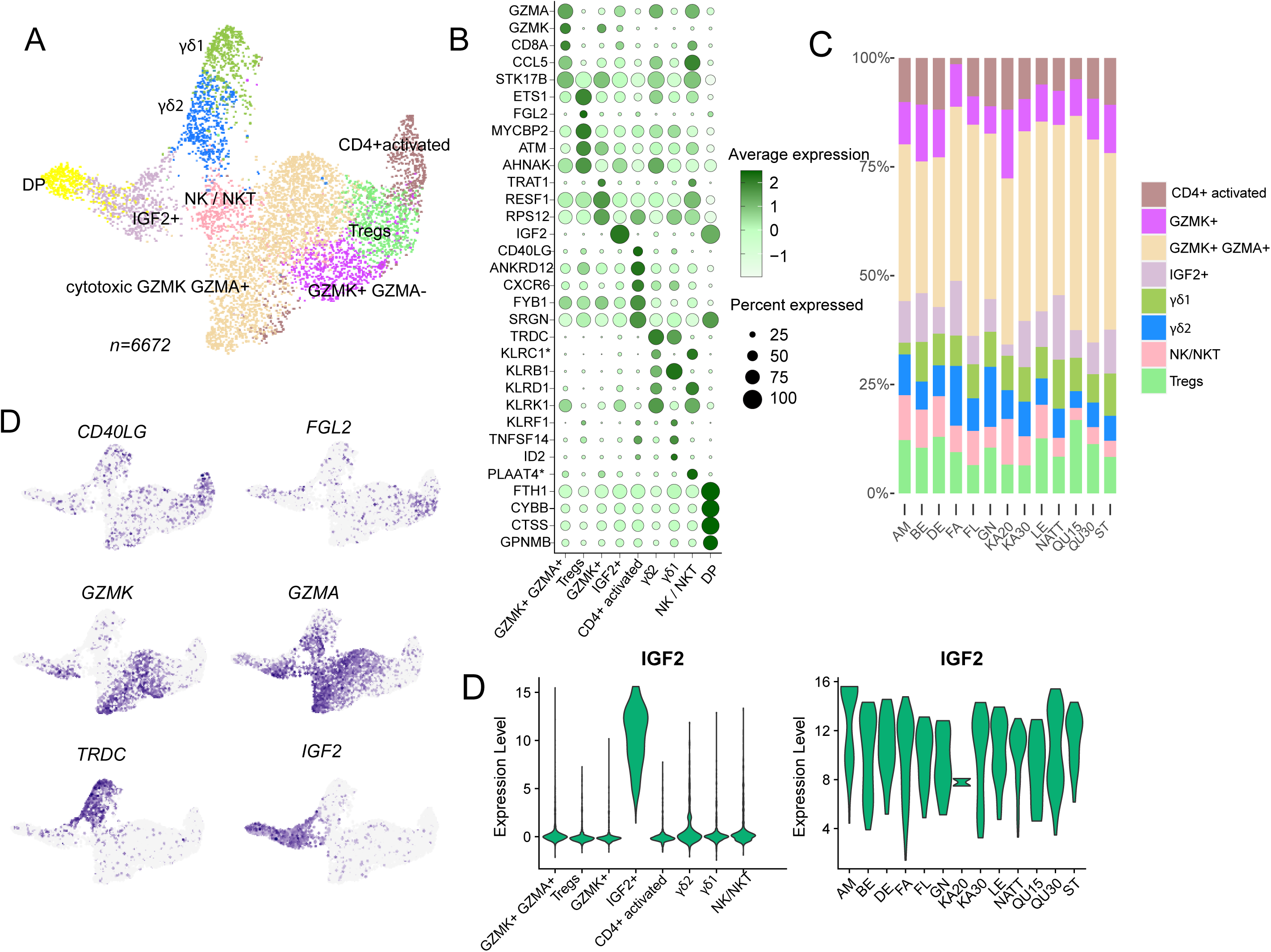
Re-clustering of T cells. A) Re-clustering of T cell population revealed several subsets: cytotoxic T cells (*GZMA, GZMA, CCL5*), γδ T subsets (*TRDC),* IGF2^+^ T cells (*IGF2*), putative Tregs (*FGL2, ETS1*), activated CD4^+^ T cells (*CD40LG*). NK/NKT cells (*KLRD1, KLRK1*). A small subset of cells expressing macrophage markers clustered separately (DP). B) Bubble plots showing expression of top cluster markers. C) Composition plots showing proportion of T cell subsets in individual horses. D) Selected feature expression mapped onto the UMAP plot. E) Insulin-like growth factor 2 (*IGF2)* was selectively expressed in a subset of T cell (left violin panel). Expression of *IFG2* per horse in the IGF2^+^ T cell cluster (right violin panel).

**Figure 7.**
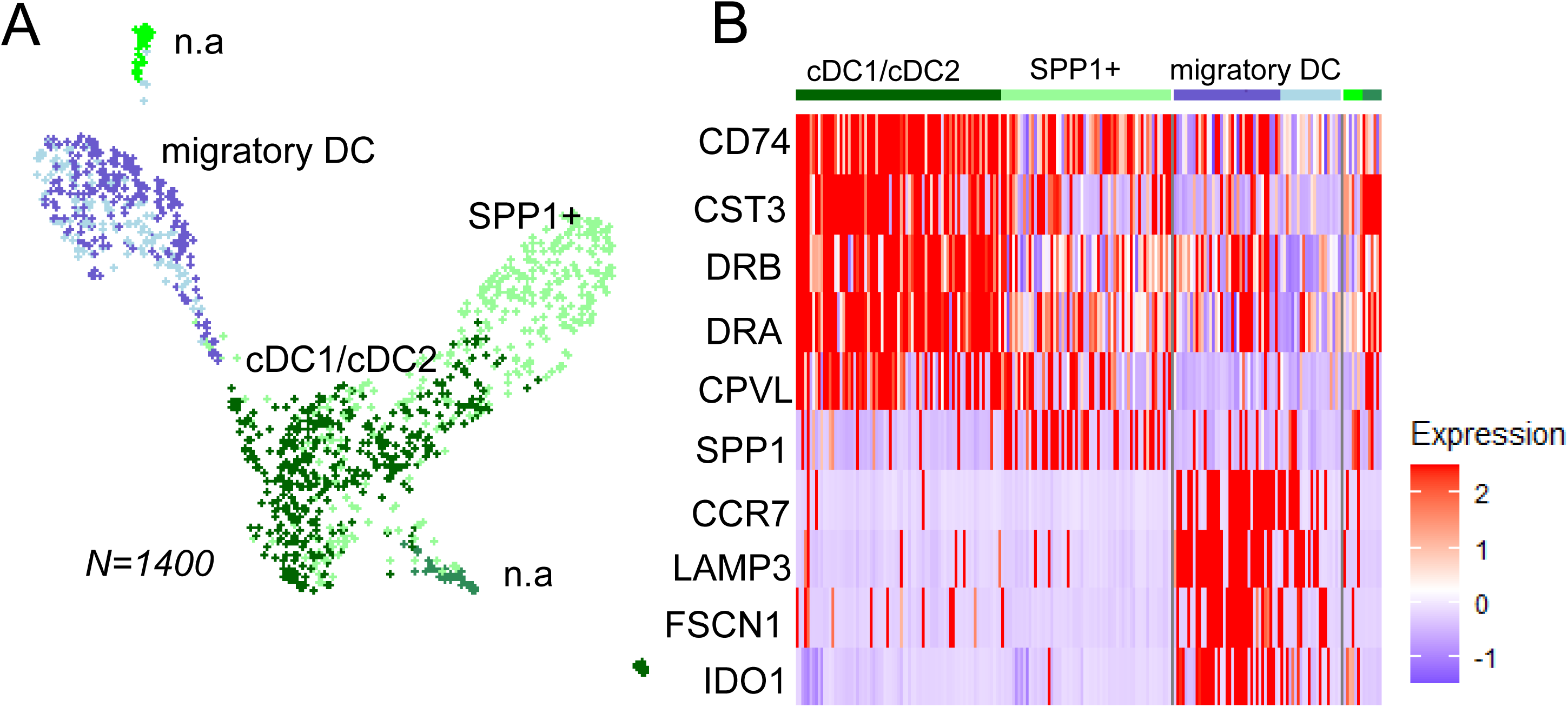
Reclustering of dendritic cells. A) UMAP visualization showing independent re-clustering of the dendritic cells. B) Conventional DCs were marked by e.g *CD74* and *CST3* expression, Migratory/activated DCs specifically expressed e.g *CCR7* and *LAMP3*. A population of DCs expressed higher *SPP1* (osteopontin) which is considered a marker of inflammation.

Cytotoxic CD8^+^ T cells were assigned based on high expression of granzymes *GZMK, GZMA*, as well as *CCL5* and the presence of *CD8A*. Moreover, a subset of T cells exhibiting moderate *GZMK* but lower *GZMA* (*GZMK*^+,^ *GZMA*^low^) were observed. T cells subsets expressing *GZMK* but lower *GZMA* has been observed in equine peripheral blood (4) but an exact annotation remain unclear. Notably, a group of T cells exhibited selectively high expression of insulin-like growth factor 2 (*IGF2*), along with a moderate cytotoxic profile (*GZMA, CCL5*). Although a relatively small number of cells were in total (n=576), all horses were represented in the *IGF2*^+^ cluster (Figure 6). *IGF2* was also among the genes detected in T cells in a scRNA-seq study of BAL cells from severe equine asthma and healthy controls (2). A role for the IGF system has been implicated in human asthma (30–32), in the regulation of Th17 cells in human autoimmune diseases (33), as well as in fibrosis (34). However, to our knowledge, the specific role of *IGF2* expression in T cells has not been previously discussed. Further studies will be required to elucidate the potential significance of this finding in the context of equine asthma.

The cluster labeled NK/NKT was characterized by the expression of killer cell lectin-like receptor genes (*KLRD1, KLRK1, KLRC1*). This cluster likely comprises either NK-like T cells or NK cells, or is a mix of both cell types. Considerable overlap in transcriptional profiles can make these cell populations challenging to annotate (35). However, similar cell populations have previously been found in equine BAL and were annotated as NK-like T cells (NKT) (2).

Two clusters were identified as γδ T cells based on high expression of the T cell receptor delta gene (*TRDC*). One subset (denoted here as γδ2) was characterized by higher cytotoxic potential, as indicated by elevated expression of *GZMA, KLRD1*, *KLRC1*, and *KLRK1*. The other subset (denoted here as γδ1) did not exhibit a strong cytotoxic profile but expressed higher levels of *KLRB1*. In humans, γδ T cells are subclassified based on TCR δ chain usage, such as Vδ1 and Vδ2. These subsets are primarily found in mucosal environments and peripheral blood, respectively and is proposed to have distinct roles in tissue homeostasis and inflammation (36,37). It is likely that similar subsets of γδ T cells exist in equines. However, since TCR δ variable (TRDV) gene usage was not analyzed in this study, it remains undetermined whether the two γδ T cell populations identified in equine BAL correspond to any previously characterized human γδ T cell subsets.

Additionally, two T cell clusters were absent of cytotoxic genes and CD8 expression. Based on differential expression of CD154 (*CD40LG*), one of these populations may constitute activated CD4^+^ T cells (38). The second population was labeled as putative Tregs. Although there was no direct support for this annotation by detection of classical Treg markers such as *FOXP3, IL2RA*, and *CTLA4*, there was upregulation of other genes that have been previously implicated in Tregs (*ETS1, FGL2*) (39,40). However, we were not able to selectively identify naïve T cells as were previously found in equine BAL(2), and as CD4^+^ and classical Treg marker expression were low in the HIVE T cell data, these cell labels are to be considered preliminary.

In summary, a moderate number of T cells were re-clustered, which may have constrained the depth of the analysis and limited the identification of specific T helper subsets. Nevertheless, several of the T cell types previously identified in equine BAL (1,2) were also found in the HIVE data. Additionally, although previous studies have found γδ T cells in BAL, two distinct γδ T cell populations were detected in the HIVE data, along with a population of T cells with notably high *IGF2* expression. However, the annotations of putative CD4^+^ populations in the HIVE data remain ambiguous. B-cells were not detected, in contrasts with previous studies of EA using 10x Genomics’ methods.

### Dendritic cells, neutrophils and mast cells as detected by the HIVE method

#### Dendritic cells

The dendritic cells were re-clustered into subtypes that have been previously observed in both equines and humans. Conventional dendritic cells (cDCs) were identified based on the expression of *CD74, CST3, DRA*, and *CPVL* (41). However, in this integrated dendritic cell data subset, we were unable to distinguish the conventional DCs into cDC1 and cDC2 subgroups, as was possible in previous studies (1,2). A group of *SPP1^+^* DCs was identified, potentially representing an inflammatory subset (42). Migratory/activated DCs, marked by *CCR7, LAMP3*, and *IDO1*, respectively was also observed, consistent with previous findings in equine BAL (1,2). Additionally, two very small clusters were present within the dendritic cell subset, but we were unable to annotate those.

#### Neutrophils and mast cells

Four subsets were observed upon re-clustering neutrophils (Figure 8, S5 Data), consistent with other studies reporting neutrophil heterogeneity (43). Among these, the previously observed interferon-stimulated (ISG^High^) neutrophils were also detected (1,2). A previous study by Sage et al(2). observed specific asthma signatures in neutrophils from horses with neutrophilic sEA, including the upregulation of *CHI3L1* and *MAPK13*. These genes were not detected in the neutrophil subsets in the current study. However, the majority of the samples analyzed here were from a different phenotype, mastocytic mEA. Of particular note, a small cluster exhibited a distinct profile characterized by the expression of *ADAMDEC1*, *GAPT*, G*LS*, *SYNE1*, *DHRS7*, and the eotaxin receptor *CCR3* (44) (Figure 8 and S5 Data). These cells were labelled as eosinophils.

**Figure 8.**
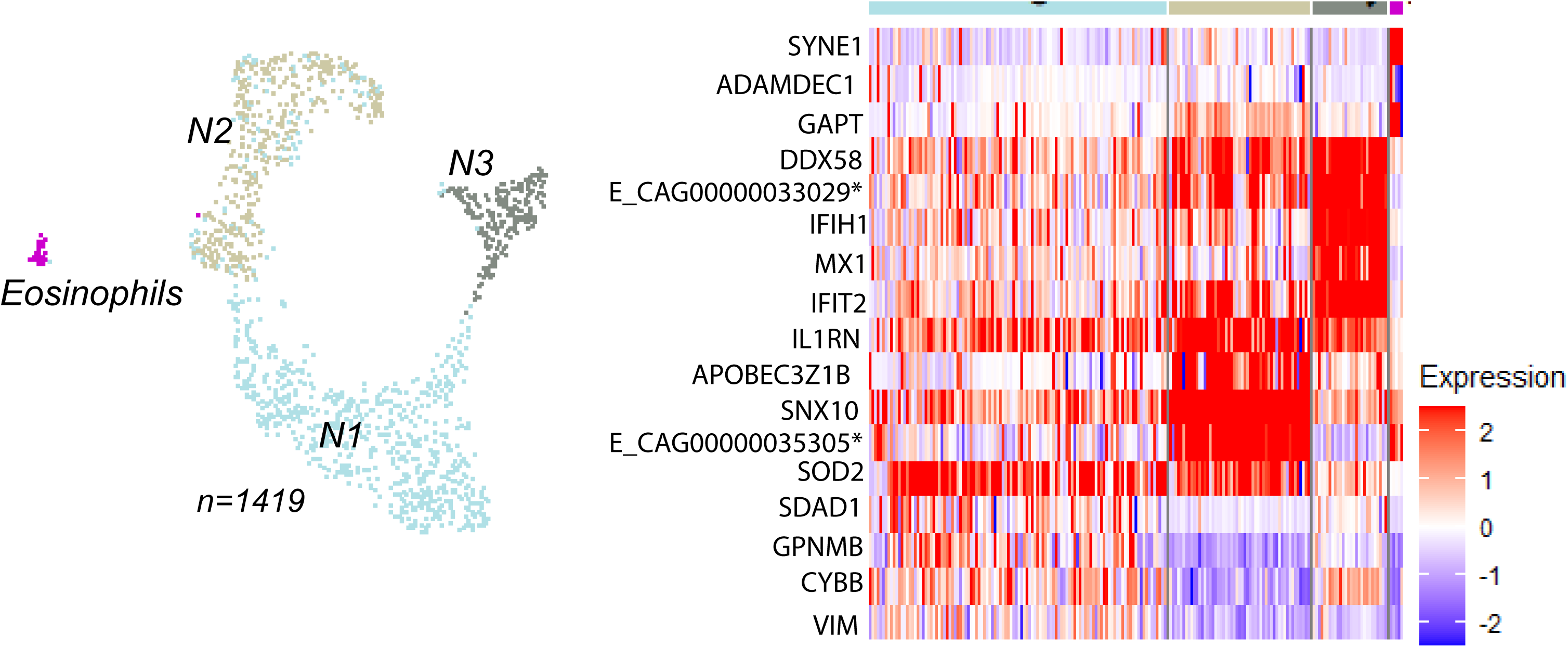
Re-clustering of neutrophils. A). UMAP visualization showing independent re-clustering of the neutrophil population. Four subsets of cell were identified. B). Clusters N1 and N2 had similar feature expression, though N2 exhibited higher expression of *SNX10* and A*POBEC3Z1B.* Cluster N3 was characterized by elevated expression of interferon-stimulated genes (ISG). Additionally, a small cluster of eosinophils displayed a very distinct profile, marked by the expression of e.g *SYNE1, ADAMDEC1*, and *GAPT*.

The small number of mast cells analyzed within the total BAL cell population were quite homogeneous, though a subset showed a subtle increase in the expression of macrophage-associated genes (Figure 9). Another small subset exhibited higher IGF2 expression. A previous study that included cells from mastocytic horses and healthy controls demonstrated a small subset of mast cells characterized by elevated *FKBP5* and *LTC4S* expression, which was primarily derived from asthma cases rather than control horses (1). However, healthy controls were not included in this evaluation so this finding could not be corroborated. It is also worth noting that a study comparing neutrophilic severe asthma cases with healthy controls found the mast cell population to be highly homogeneous, regardless of disease status (2).

**Figure 9.**
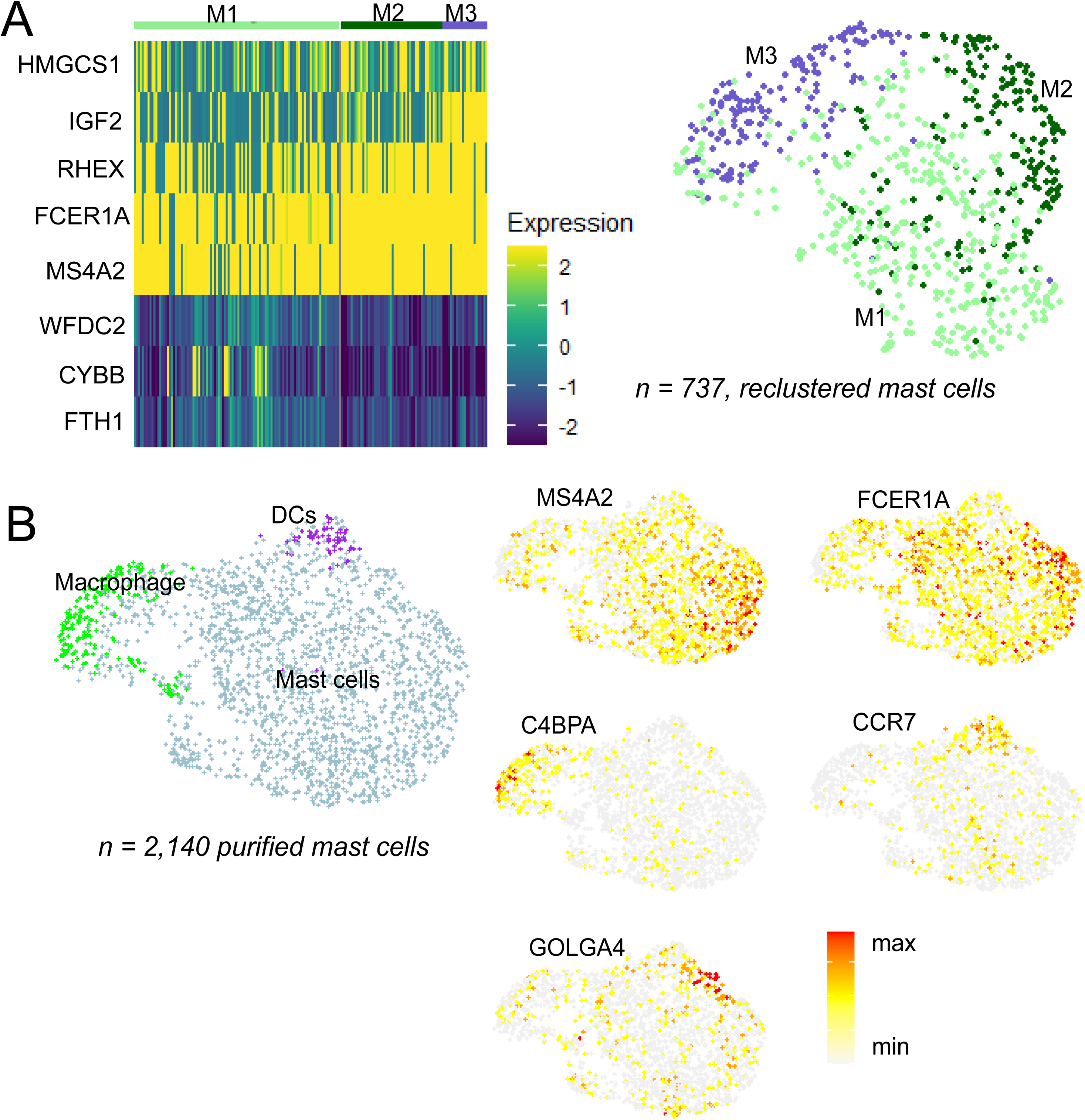
Analysis of mast cells in BAL. A) Heatmap showing the expression of top differentially expressed genes across three clusters observed when re-clustering 737 mast cells from 13 BAL samples (clusters M1-M3 in the UMAP). B) Clustering of 2,140 purified mast cells revealed mainly mast cells (90%) but also small populations of macrophages and dendritic cells F) Feature plots illustrating expression of selected genes in the purified mast cells: general mast cell population marker (*MS4A2, FCER1A*), alveolar macrophage marker (*C4BPA*), dendritic cell marker (*CCR7*) and the Golgi-associated gene *GOLGA4*.

Due to the low recapitulation of mast cells in the total BAL samples, we wanted to further assess the HIVE method’s capability to capture mast cells and identify potential loss points in the workflow. To this end, we performed HIVE scRNA-seq on a sample of purified equine mast cells isolated from a single BAL sample (library #14, sample ID = UF). The library was generated using the same procedure as for the initial BAL samples, including the same cell dilution buffer. The analysis resulted in 2,140 cells, with 90% identified as mast cells based on the expression of classical mast cell markers (*MS4A2*, *FCER1A*) (Figure 9). A subset of mast cells was observed to express high levels of mRNA encoding the Golgi apparatus-associated proteins *GOLGA4*, involved in vesicular trafficking in the Golgi apparatus. Additionally, small subsets of dendritic cells (*CCR7*, *LAMP3*) and alveolar macrophages (*MARCO*, *C4BPA*) were identified, (Figure 9, S5 Data).

In summary, while the recapitulation of neutrophils and mast cells in total BAL samples were low, the HIVE method captured expected numbers of purified mast cell. A small subset of eosinophil was detected which has not previously been characterized in equine BAL.

### Characteristics of the HIVE method in comparison with alternative scRNA-seq methods

While droplet fluidics-based methods, like those provided by 10x Genomics, have been the most widely used for scRNA-seq, the range of available techniques is rapidly expanding.

Various approaches, which do not require specialized instruments, have emerged, offering greater flexibility and potentially reduced costs. For example, products from Parse and Scale Biosciences utilize approaches for split-pool combinatorial barcoding in protocols that involve early cell fixation while also allowing for high sample multiplexing (45,46). Another notable method is PipSeq (formerly from Fluent-Bio, now under Illumina Inc.), which employs a vortex-based emulsification technique to partition cells along with barcoded particles (47). The primary advantage of the HIVE method, compared to other methods, is its convenient cell handling procedure, which can be performed directly at the sampling site without the need for additional equipment beyond pipettes.

A significant limitation of this method evaluation is the absence of a parallel comparison of quality metrics and sensitivity (e.g., gene detection levels) of the HIVE method with alternative methods using the same BAL samples. In the absence of such comparisons, we compiled metrics that indicate a method’s sensitivity (i.e., gene and UMI metrics) from previously published studies using related sample types (Table 4). The HIVE method demonstrated higher sensitivity than Drop-seq data but detected fewer genes and UMIs compared to 10X Genomics.

**Table 4.** Metrics comparison across studies and different scRNA-seq protocols.

| Method | Study | Sample type | N samples | genes/cell range | genes/cell median | UMIs/cell median |
| --- | --- | --- | --- | --- | --- | --- |
| HIVE | Present study | Horse/BAL | 13 | 492-1189 | 673 | 1055 |
| Drop-seq | Riihimäki *et al* (1) | Horse/BAL | 11 | 410-748 | 602 | 931 |
| Chromium 3'/v.3 | Sage *et al* (2) | Horse/BAL | 12 | 589-1351 | 778 | 1856 |
| Chromium 5' | Feys *et al* (48) | Human/BAL | 43 | 218-1510 | 754 | 2223 |
| Chromium 3'/v.2 | Patel *et al* (4) | Horse/PBMC | 7 | 881-996<sup>a</sup> | 962 | n.a<sup>b</sup> |

<sup>a</sup>corresponding numbers pre-ESAT correction were = 560-607 genes/cell, <sup>b</sup>n.a= data were not available

In a recent study that conducted a comprehensive evaluation of scRNA kits, the HIVE method demonstrated a moderate-to-low level of gene and UMI recovery, placing it in the lower performance tier compared to top-performing methods such as 10x Genomics (25). Additionally, the HIVE method did not perform as well in accurately replicating cell type distribution and discriminating cell clusters compared to methods like 10x Genomics. It showed larger deviations from the reference cell type proportions determined by CyTOF, particularly in the distribution of monocytes and T cells (25), which is also consistent with our observations in this study. A second study that compared HIVE, Parse, and 10X Genomics Flex confirmed discrepancies in cell type distributions when compared to flow cytometry, but nonetheless reported that all three methods generated high-quality data from blood neutrophils (49).

The HIVE v1 protocol exhibited considerable variability in the number of recovered cells across the samples in our study, ranging from 1,000 to 7,000 cells. We speculate that this variability might partly stem from a specific step in the library preparation protocol, where the beads from the HIVE collector (containing the captured cell transcriptomes) are centrifuged and transferred to plate wells. This bead pellet is easily disturbed, which can lead to a suboptimal number of beads (and hence transcriptomes) being carried over to downstream processing. Another common cause of discrepancies between targeted and recovered number of cells in scRNA-seq experiments is incorrect quantification of the cell suspensions. It should be noted that in this study the first version of the kit (HIVE v1) were used, but the most recent version (HIVE CLX) allows for a broader range of cell loading (500–60,000 cells/HIVE), potentially enabling higher cell recovery.

The low recovery of neutrophils and mast cells in total BAL our study was unexpected, especially since we anticipated better representation of these cells with the HIVE method in comparison to droplet-based methods. This was particularly puzzling given that several BAL samples included in the study contained 10-20% mast cells and one sample contained 30% neutrophils according to cytology. Less stringent data filtering did not improve the recovery of either neutrophils or mast cells in the total BAL samples (data not shown). In contrast, loading purified mast cells onto the HIVE collector yielded an expected number of recovered mast cells. One possibility is that the purified mast cells represent terminally differentiated, tissue-resident mast cells, which have been shown to be long-lived (50) and that the mast cells lost during the BAL cell preparation were either immature or activated mast cells. Therefore, the apparent loss of granulocytes and mast cells observed when the HIVE method (and other scRNA-seq methods) is applied to total BAL cells is likely not due to the method itself, but rather to factors in the BAL sample processing workflow.

## Conclusion

Despite deviations from the expected cell-type distributions in cytology, the HIVE data identified cell types that were generally comparable to those in previous studies. While the HIVE method may not match the sensitivity of methods from10X Genomics’ methods, its practical and rapid sample handling procedure makes it a useful option for both field and clinical site sampling. However, our findings suggest that further optimization of sample pre-processing and workflow is needed to improve the recovery of equine BAL granulocytes and achieve more accurate assessment of cell-type composition.

## Data availability

Raw sequencing data will be deposited at NCBI SRA with accession nr: PRJNA1085264. The R-scripts and processed data used for the analysis, as well as clinical information will be found at: link DOI: 10.17044/scilifelab.25471861.

## Acknowledgements

Sequencing was performed by the SciLifeLab National Genomics Infrastructure (NGI) SNP&SEQ unit at Uppsala University. NGI is funded by SciLifeLab, the Knut and Alice Wallenberg Foundation, and the Swedish Research Council. The computation analyses were enabled by resources provided by the National Academic Infrastructure for Supercomputing in Sweden (NAISS) at UPPMAX, funded by the Swedish Research Council. We thank Anders Lundmark for assistance in running the BeeNet software.

## Author contributions

KF performed experiments and analyzed data, MR performed clinical examinations and BAL samplings. JN provided resources and expertise in bioinformatics, SA performed experiments, SW provided expertise in immunology. AR conceived the study, analyzed the data and wrote the manuscript. All authors read and edited the manuscript.

## Supplementary Files

S1. Figure, Example of TapeStation profiles for HIVE libraries. S2. Figure, QC plots.

S1. Table, Age, sex, breed and sampling dates.

S2. Table, Cytology data for Drop-seq samples.

S1. Data, BeeNet metrics

S2. Data, Cluster and major cell type marker genes for initial HIVE data

S3. Data, Cell type marker genes for the sample handling test data

S4. Data, Cell type marker genes for mastocytic HIVE and DropSeq data.

S5. Data, Marker genes from independent re-clustering of cell types and isolated mast cells

## References

1. Riihimäki M, Fegraeus K, Nordlund J, Waern I, Wernersson S, Akula S, et al. Single-cell transcriptomics delineates the immune cell landscape in equine lower airways and reveals upregulation of FKBP5 in horses with asthma. Sci Rep. 2023 Sep;13(1):16261.

2. Sage SE, Leeb T, Jagannathan V, Gerber V. Single-cell profiling of bronchoalveolar cells reveals a Th17 signature in neutrophilic severe equine asthma. Immunology. 2023 Dec;

3. Sage SE, Nicholson P, Peters LM, Leeb T, Jagannathan V, Gerber V. Single-cell gene expression analysis of cryopreserved equine bronchoalveolar cells. Front Immunol. 2022;13:929922.

4. Patel RS, Tomlinson JE, Divers TJ, Van de Walle GR, Rosenberg BR. Single-cell resolution landscape of equine peripheral blood mononuclear cells reveals diverse cell types including T-bet+ B cells. BMC Biol. 2021 Jan;19(1):13.

5. Deeg CA, Hauck SM, Amann B, Pompetzki D, Altmann F, Raith A, et al. Equine recurrent uveitis–a spontaneous horse model of uveitis. Ophthalmic Res. 2008;40(3– 4):151–3.

6. Bullone M, Lavoie JP. The equine asthma model of airway remodeling: from a veterinary to a human perspective. Cell Tissue Res. 2020 May;380(2):223–36.

7. Bullone M, Lavoie JP. Asthma ‘of horses and men’–how can equine heaves help us better understand human asthma immunopathology and its functional consequences? Mol Immunol. 2015 Jul;66(1):97–105.

8. Jensen-Jarolim E, Einhorn L, Herrmann I, Thalhammer JG, Panakova L. Pollen Allergies in Humans and their Dogs, Cats and Horses: Differences and Similarities. Clin Transl Allergy. 2015;5:15.

9. Ellis KL, Contino EK, Nout-Lomas YS. Poor performance in the horse: Diagnosing the non-orthopaedic causes. Equine Vet Educ. 2023;35(4):208–24.

10. Bond S, Léguillette R, Richard EA, Couetil L, Lavoie JP, Martin JG, et al. Equine asthma: Integrative biologic relevance of a recently proposed nomenclature. J Vet Intern Med. 2018 Nov;32(6):2088–98.

11. Couetil L, Cardwell JM, Leguillette R, Mazan M, Richard E, Bienzle D, et al. Equine Asthma: Current Understanding and Future Directions. Front Vet Sci. 2020;7:450.

12. Kinnison T, McGilvray TA, Couëtil LL, Smith KC, Wylie CE, Bacigalupo SA, et al. Mild-moderate equine asthma: A scoping review of evidence supporting the consensus definition. Vet J Lond Engl 1997. 2022 Aug;286:105865.

13. Cian F, Monti P, Durham A. Cytology of the lower respiratory tract in horses: An updated review. Equine Vet Educ. 2015 Oct;27(10):544–53.

14. Airway Diagnostics: Bronchoalveolar Lavage, Tracheal Wash, and Pleural Fluid - Available from: https://pubmed.ncbi.nlm.nih.gov/32145836/

15. Baratchi S, Danish H, Chheang C, Zhou Y, Huang A, Lai A, et al. Piezo1 expression in neutrophils regulates shear-induced NETosis. Nat Commun. 2024 Aug 22;15(1):7023.

16. Ledderose C, Hashiguchi N, Valsami EA, Rusu C, Junger WG. Optimized flow cytometry assays to monitor neutrophil activation in human and mouse whole blood samples. J Immunol Methods. 2023 Jan;512:113403.

17. Gierahn TM, Wadsworth MH, Hughes TK, Bryson BD, Butler A, Satija R, et al. Seq-Well: portable, low-cost RNA sequencing of single cells at high throughput. Nat Methods. 2017 Apr;14(4):395–8.

18. Akula S, Riihimäki M, Waern I, Åbrink M, Raine A, Hellman L, et al. Quantitative Transcriptome Analysis of Purified Equine Mast Cells Identifies a Dominant Mucosal Mast Cell Population with Possible Inflammatory Functions in Airways of Asthmatic Horses. Int J Mol Sci. 2022 Nov;23(22):13976.

19. Hafemeister C, Satija R. Normalization and variance stabilization of single-cell RNA-seq data using regularized negative binomial regression. Genome Biol. 2019 Dec;20(1):296.

20. McGinnis CS, Murrow LM, Gartner ZJ. DoubletFinder: Doublet Detection in Single-Cell RNA Sequencing Data Using Artificial Nearest Neighbors. Cell Syst. 2019 Apr;8(4):329–337.e4.

21. Zappia L, Oshlack A. Clustering trees: a visualization for evaluating clusterings at multiple resolutions. GigaScience 2018 Jul 1;7(7).

22. Trivedi NN, Tamraz B, Chu C, Kwok PY, Caughey GH. Human subjects are protected from mast cell tryptase deficiency despite frequent inheritance of loss-of-function mutations. J Allergy Clin Immunol. 2009 Nov;124(5):1099–1105.e4.

23. participants in the 1st Human Cell Atlas Jamboree, Lun ATL, Riesenfeld S, Andrews T, Dao TP, Gomes T, et al. EmptyDrops: distinguishing cells from empty droplets in droplet-based single-cell RNA sequencing data. Genome Biol. 2019 Dec;20(1):63.

24. Uhlén M, Fagerberg L, Hallström BM, Lindskog C, Oksvold P, Mardinoglu A, et al. Tissue-based map of the human proteome. Science. 2015 Jan 23;347(6220):1260419.

25. De Simone M, Hoover J, Lau J, Bennet H, Wu B, Chen C, et al. Comparative Analysis of Commercial Single-Cell RNA Sequencing Technologies [Internet]. 2024 [cited 2024 Sep 20]. Available from: http://biorxiv.org/lookup/doi/10.1101/2024.06.18.599579

26. Smith TD, Tse MJ, Read EL, Liu WF. Regulation of macrophage polarization and plasticity by complex activation signals. Integr Biol. 2016 Sep 12;8(9):946–55.

27. Lenga Ma Bonda W, Fournet M, Zhai R, Lutz J, Blondonnet R, Bourgne C, et al. Receptor for Advanced Glycation End-Products Promotes Activation of Alveolar Macrophages through the NLRP3 Inflammasome/TXNIP Axis in Acute Lung Injury. Int J Mol Sci. 2022 Oct 1;23(19):11659.

28. Theobald H, Bejarano DA, Katzmarski N, Haub J, Schulte-Schrepping J, Yu J, et al. Apolipoprotein E controls Dectin-1-dependent development of monocyte-derived alveolar macrophages upon pulmonary β-glucan-induced inflammatory adaptation. Nat Immunol. 2024 Jun;25(6):994–1006.

29. Kang H, Bienzle D, Lee GKC, Piché É, Viel L, Odemuyiwa SO, et al. Flow cytometric analysis of equine bronchoalveolar lavage fluid cells in horses with and without severe equine asthma. Vet Pathol. 2022 Jan;59(1):91–9.

30. He J, Mu M, Wang H, Ma H, Tang X, Fang Q, et al. Upregulated IGF-1 in the lungs of asthmatic mice originates from alveolar macrophages. Mol Med Rep. 2019 Feb;19(2):1266–71.

31. Vázquez-Mera S, Pichel JG, Salgado FJ. Involvement of IGF Proteins in Severe Allergic Asthma: New Roles for Old Players. Arch Bronconeumol. 2021 Dec;57(12):731–2.

32. Yang G, Geng XR, Song JP, Wu Y, Yan H, Zhan Z, et al. Insulin-like growth factor 2 enhances regulatory T-cell functions and suppresses food allergy in an experimental model. J Allergy Clin Immunol. 2014 Jun;133(6):1702–1708.e5.

33. DiToro D, Harbour SN, Bando JK, Benavides G, Witte S, Laufer VA, et al. Insulin-Like Growth Factors Are Key Regulators of T Helper 17 Regulatory T Cell Balance in Autoimmunity. Immunity. 2020 Apr;52(4):650–667.e10.

34. Zhu Y, Chen L, Song B, Cui Z, Chen G, Yu Z, et al. Insulin-like Growth Factor-2 (IGF-2) in Fibrosis. Biomolecules. 2022 Oct 25;12(11):1557.

35. ImmGen Project Consortium, Cohen NR, Brennan PJ, Shay T, Watts GF, Brigl M, et al. Shared and distinct transcriptional programs underlie the hybrid nature of iNKT cells. Nat Immunol. 2013 Jan;14(1):90–9.

36. Wu D, Wu P, Qiu F, Wei Q, Huang J. Human γδT-cell subsets and their involvement in tumor immunity. Cell Mol Immunol. 2017 Mar;14(3):245–53.

37. Chen Y, Li J, Zeng X, Yuan W, Xu Y. γδ T cells and their roles in immunotherapy: a narrative review. Ann Blood. 2022 Dec;7:42–42.

38. Grewal IS, Flavell RA. The Role of CD40 Ligand in Costimulation and T-Cell Activation. Immunol Rev. 1996 Oct;153(1):85–106.

39. Hou XX, Wang XQ, Zhou WJ, Li DJ. Regulatory T cells induce polarization of pro-repair macrophages by secreting sFGL2 into the endometriotic milieu. Commun Biol. 2021 Apr 23;4(1):499.

40. Wang J, Vuitton DA, Müller N, Hemphill A, Spiliotis M, Blagosklonov O, et al. Deletion of Fibrinogen-like Protein 2 (FGL-2), a Novel CD4+ CD25+ Treg Effector Molecule, Leads to Improved Control of Echinococcus multilocularis Infection in Mice. Flisser A, editor. PLoS Negl Trop Dis. 2015 May 8;9(5):e0003755.

41. Naessens T, Morias Y, Hamrud E, Gehrmann U, Budida R, Mattsson J, et al. Human Lung Conventional Dendritic Cells Orchestrate Lymphoid Neogenesis during Chronic Obstructive Pulmonary Disease. Am J Respir Crit Care Med. 2020 Aug 15;202(4):535– 48.

42. Tang S, Hu H, Li M, Zhang K, Wu Q, Liu X, et al. OPN promotes pro-inflammatory cytokine expression via ERK/JNK pathway and M1 macrophage polarization in Rosacea. Front Immunol. 2024 Jan 5;14:1285951.

43. Silvestre-Roig C, Fridlender ZG, Glogauer M, Scapini P. Neutrophil Diversity in Health and Disease. Trends Immunol. 2019 Jul;40(7):565–83.

44. Oliveira SHP, Lukacs NW. The role of chemokines and chemokine receptors in eosinophil activation during inflammatory allergic reactions. Braz J Med Biol Res. 2003 Nov;36(11):1455–63.

45. Rosenberg AB, Roco CM, Muscat RA, Kuchina A, Sample P, Yao Z, et al. Single-cell profiling of the developing mouse brain and spinal cord with split-pool barcoding. Science. 2018 Apr 13;360(6385):176–82.

46. Mulqueen RM, Pokholok D, O’Connell BL, Thornton CA, Zhang F, O’Roak BJ, et al. High-content single-cell combinatorial indexing. Nat Biotechnol. 2021 Dec;39(12):1574– 80.

47. Clark IC, Fontanez KM, Meltzer RH, Xue Y, Hayford C, May-Zhang A, et al. Microfluidics-free single-cell genomics with templated emulsification. Nat Biotechnol. 2023 Nov;41(11):1557–66.

48. Feys S, Carvalho A, Clancy CJ, Gangneux JP, Hoenigl M, Lagrou K, et al. Influenza-associated and COVID-19-associated pulmonary aspergillosis in critically ill patients. Lancet Respir Med. 2024 Sep;12(9):728–42.

49. Hatje K, Schneider K, Danilin S, Koechl F, Giroud N, Juglair L, et al. Comparison of Fixed Single Cell RNA-seq Methods to Enable Transcriptome Profiling of Neutrophils in Clinical Samples. http://biorxiv.org/lookup/doi/10.1101/2024.08.14.607767

50. Gilfillan AM, Austin SJ, Metcalfe DD. Mast Cell Biology: Introduction and Overview. In: Gilfillan AM, Metcalfe DD, editors. Mast Cell Biology. Boston, MA: Springer US; 2011. p. 2–12. (Advances in Experimental Medicine and Biology; vol. 716).

