## Supplementary figures and images for "Exploring a pico-well based scRNA-seq method (HIVE) for simplified processing of equine bronchoalveolar lavage cells"

### S1 Figure

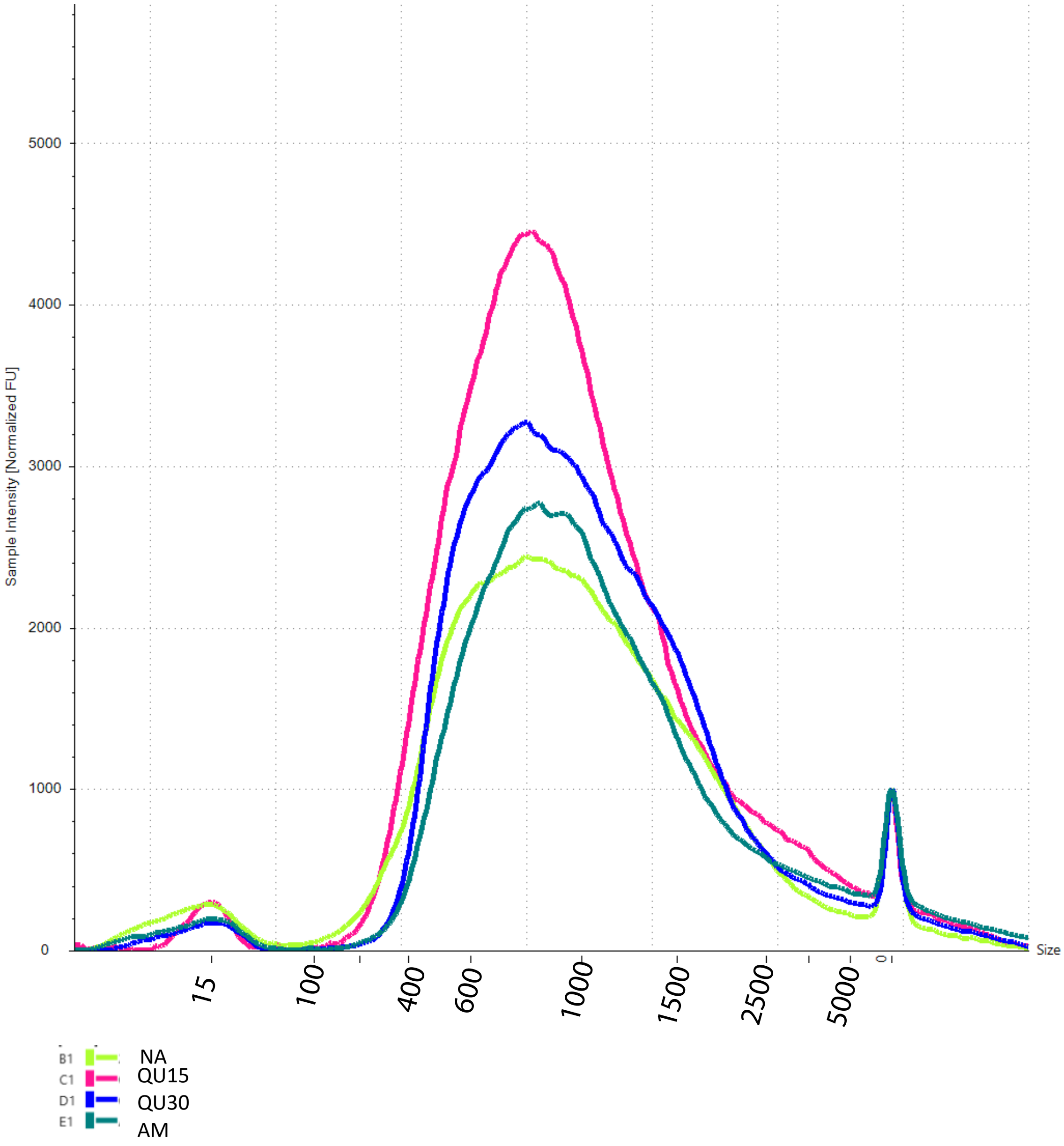


**S1 Figure**. Example of TapeStation size profiles for HIVE libraries
