## Supplementary material for "Exploring a pico-well based scRNA-seq method (HIVE) for simplified processing of equine bronchoalveolar lavage cells": S2 Figure

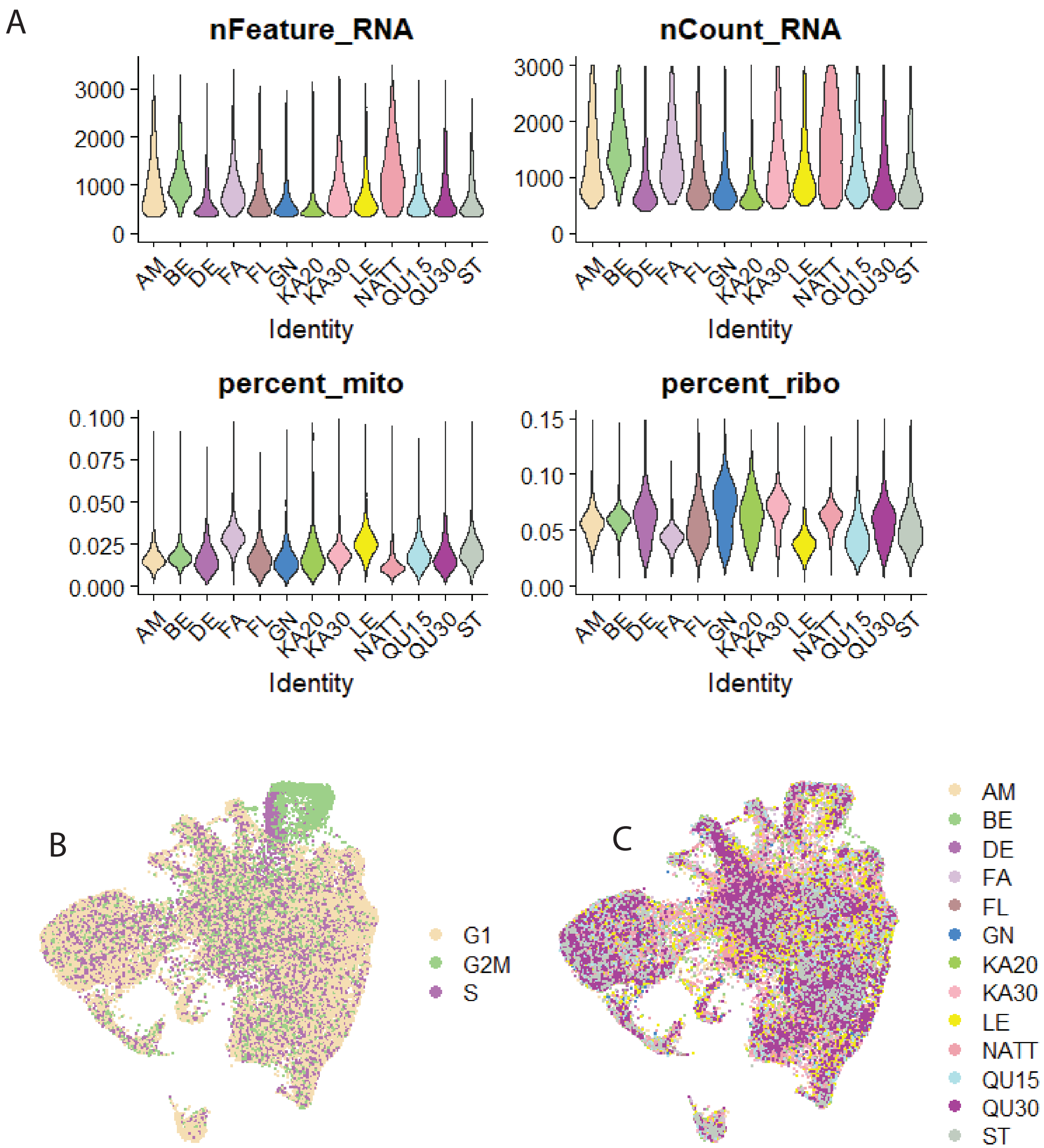


**S2 Figure.**
A. Violin plots showing QC metrics for 13 HIVE libraries. Genes (nFeatures)/cell. UMIs (nCounts)/cell. Percent mitochondrial and ribosomal counts/cell.
B. UMAP plot showing cells colored by cell cycle scores.
C. UMAP plot showing cells color by sample identity.
