## Supplementary material for "Exploring a pico-well based scRNA-seq method (HIVE) for simplified processing of equine bronchoalveolar lavage cells": S1 Table

**S1 Table. Sample and batch information**

| Horse ID | Age | Sex | Breed | Sample date | Library prep  batch | Sequencing  batch* |
| --- | --- | --- | --- | --- | --- | --- |
| NA | 5 | M | Icelandic horse | 2021-11-08 | 1 | 1 |
| DE | 13 | M | Danish Warmblood | 2022-02-07 | 2 | 1 |
| QU | 8 | M | Swedish Warmblood | 2022-03-14 | 2 | 1 |
| AM | 18 | M | Swedish Warmblood | 2022-03-21 | 2 | 1 |
| ST | na* | G | Welsh pony | 2022-05-03 | 3 | 1 |
| BE | 12 | M | Swedish Warmblood | 2022-05-02 | 4 | 1 |
| FA | 8 | M | Shetland pony | 2022-05-16 | 4 | 1 |
| LE | 13 | G | Swedish Warmblood | 2022-05-23 | 4 | 1 |
| KA | 16 | G | Icelandic horse | 2022-06-13 | 5 | 1 |
| GN | 20 | G | Icelandic horse | 2022-09-05 | 5 | 1 |
| FL | 15 | M | Swedish Warmblood | 2022-09-26 | 5 | 1 |
| UF | 7 | G | Swedish Warmblood | 2021-09-27 | 7 | 2 |
| AT | 16 | M | Swedish Warmblood | 2023-08-28 | 6 | 3 |
| TT | 15 | G | Swedish Warmblood | 2023-11-09 | 6 | 3 |

*not available

*horses NA, DE, QU, AM, ST, BE, FA, LE, KA, GN, FL were sequenced together on a NovaSeq 6000 S4 flow cell. Libraries from horse AT and TT were sequenced together on a NovaSeq 6000 SP flow cell. Horse UF were sequenced on a separate NovaSeq SP flow cell.
