## Supplementary material for "Exploring a pico-well based scRNA-seq method (HIVE) for simplified processing of equine bronchoalveolar lavage cells": S2 Table

| **ID*** | **Neutrophils**  **(%)** | **Lymphocytes**  **(%)** | **Macrophages**  **(%)** | **Eosinophils (%)** | **Mast cells**  **(%)** |
| --- | --- | --- | --- | --- | --- |
| **DS** | 1 | 35 | 52 | 1 | 11 |
| **CA** | 2 | 41 | 50 | 0 | 7 |
| **FS** | 7 | 33 | 42 | 8 | 10 |
| **NE** | 0 | 37 | 47 | 0 | 10 |
| **OD** | 4 | 39 | 28 | 18 | 9 |
| **TI** | 5 | 33 | 53 | 0 | 9 |
| **VA** | 5 | 12 | 71 | 6 | 6 |
| **VAL** | 3 | 37 | 52 | 1 | 7 |

**S2 Table. Cytology counts for horses analyzed with Drop-seq**

*Additional information regarding these horses as well as clinical scores and data availability can be found in Riihimäki et al, Scientific Reports, 2023.
